# Targeting Siglec-10/α3β1 Integrin Interactions Enhances Macrophage-Mediated Phagocytosis of Pancreatic Cancer

**DOI:** 10.1101/2025.05.06.652455

**Authors:** Pratima Saini, Gauri Mirji, S M Shamsul Islam, Lacy M. Simons, Sajad Ahmed Bhat, Amanda P. Bonfanti, Kar Muthumani, Priyesh Agrawal, Joel Cassel, Hsin-Yao Tang, Hiroaki Tateno, Rugang Zhang, Judd F. Hultquist, Rahul S. Shinde, Mohamed Abdel-Mohsen

**Author notes:** Corresponding author: Mohamed Abdel-Mohsen, Ph.D. Associate Professor, Northwestern University; Chicago, IL 60611.

## Abstract

Tumor-associated macrophages (TAMs) in the pancreatic ductal adenocarcinoma (PDAC) tumor microenvironment (TME) exhibit immunosuppressive phenotypes and impaired phagocytic activity, facilitating tumor progression and immune evasion. Here, we identify integrin α3β1, composed of ITGA3 and ITGB1 subunits, as a sialylated glycoprotein ligand for Siglec-10, an inhibitory glyco-immune checkpoint receptor highly expressed on TAMs in PDAC. The interaction between Siglec-10 on TAMs and α3β1 on PDAC cells suppresses macrophage-mediated phagocytosis, thereby promoting immune evasion. Consistently, disrupting Siglec-10 interactions with monoclonal antibodies significantly enhances macrophage phagocytosis of PDAC cells and alleviates myeloid cell-mediated inhibition of T cell proliferation and activation *in vitro*. In both a PDAC xenograft mouse model engrafted with human macrophages and a human Siglec-10 transgenic mouse model, targeting Siglec-10 with monoclonal antibodies reduces PDAC tumor growth. These findings suggest that Siglec-10 interactions are key mediators of TAM-driven immune evasion in PDAC and highlight the therapeutic potential of targeting these interactions to restore anti-tumor immunity.

**SIGNIFICANCE:** Pancreatic tumor cells exploit integrin α3β1 to engage the immunosuppressive checkpoint receptor Siglec-10 on myeloid cells, driving immune evasion, and antibody-mediated blockade of Siglec-10 restores myeloid cell–mediated anti-PDAC immunity.

## INTRODUCTION

There is an urgent need to identify novel immunological targets to improve survival rates for patients with pancreatic ductal adenocarcinoma (PDAC). This cancer remains exceptionally challenging, with a five-year survival rate of only 13% (1,2), and it shows substantial resistance to current immunotherapies (3). PDAC’s resistance to immunotherapy is thought to arise from multiple mechanisms within the tumor microenvironment (TME). Among these, tumor-associated myeloid cells, including tumor-associated macrophages (TAMs), which represent a major immune infiltrate in the TME, play a critical role in promoting treatment resistance. These myeloid cells are often reprogrammed into dysfunctional, immunosuppressive phenotypes that impair T cell effector functions and fail to phagocytose tumor cells (4–6). However, the specific factors driving this myeloid reprogramming remain poorly defined, hindering the development of effective immunotherapies for PDAC.

One emerging mechanism of myeloid immunosuppression involves alterations in tumor cell glycosylation. Advances in the field of glycobiology have revealed that many tumor cells exploit aberrant glycosylation patterns to evade immune responses (7–10). A particularly common glycomic alteration is hyper-sialylation, the process by which cancer cells upregulate sialic acid-containing glycans (sialoglycans) on their surface (11). This enables these cancer cells to bind members of the Siglec family, a group of mostly inhibitory receptors predominantly expressed on innate immune cells, including myeloid cells. Many Siglecs contain immunoreceptor tyrosine-based inhibitory motifs (ITIMs); their engagement triggers downstream inhibitory signaling cascades via SHP-1 or SHP-2 phosphorylation. These signals promote immunosuppressive phenotypes of myeloid cells (12–18). Thus, Siglecs on myeloid cells function as “glyco-immune checkpoints,” analogous to PD-1 on CD8^⁺^ T cells. Targeting the interactions between Siglecs on myeloid cells and sialic acid on cancer cells therefore holds significant potential for reprograming myeloid cells toward an immune-activated, anti-tumor state.

Multiple studies (19–24) highlight the potential of targeting Siglec-10 interactions to reprogram macrophages and restore anti-tumor immunity. For instance, sialic acid on the glycoprotein CD24, expressed by breast and ovarian cancer cells, binds to Siglec-10 on myeloid cells (20). This interaction transmits an inhibitory signal that induces myeloid cell immunosuppressive features and impairs macrophage-mediated phagocytosis of cancer cells (20). In an ovarian cancer mouse model, blockade of CD24 significantly reduced tumor growth and enhanced macrophages’ phagocytic activity (20). While the roles of other Siglecs (Siglec-7, Siglec-9, and Siglec-15) (25–31) have been explored in the context of PDAC, Siglec-10’s contribution to the immunosuppressive PDAC TME is understudied, despite its important role in other cancers (20).

In this study, we investigated the role of Siglec-10 interactions in enabling PDAC cells to evade myeloid-mediated anti-tumor activities. Unlike in ovarian and breast cancers, where CD24 is a major ligand for Siglec-10, we found that most PDAC cells do not predominantly express CD24. They do, however, bind Siglec-10, suggesting the presence of an alternative sialic acid-containing glycoprotein ligand. Using affinity pulldown assays followed by mass spectrometry, we identified integrin α3β1 (composed of ITGA3 and ITGB1 subunits) as a novel glycoprotein ligand for Siglec-10 on PDAC cells. These integrins had not previously been implicated in PDAC immune evasion. We further confirmed that ITGA3 and ITGB1 contribute to PDAC resistance to macrophage-mediated phagocytosis using *in vitro* immune-killing assays. Given that Siglec-10 on macrophages interacts with multiple glycoproteins on PDAC cells, we hypothesized that directly targeting Siglec-10 would be a more effective strategy than blocking individual ligands. To test this, we developed novel monoclonal antibodies against Siglec-10 and evaluated their ability to enhance macrophage phagocytosis of PDAC cells and to reverse myeloid cell-mediated suppression of T cell proliferation and activation. Disrupting Siglec-10 interactions with these antibodies significantly increased macrophage-mediated phagocytosis of PDAC cells and restored T cell proliferation and activation *in vitro*. Furthermore, these antibodies reduced tumor burden in both a PDAC xenograft mouse model engrafted with human macrophages and a transgenic mouse model expressing human Siglec-10.

## MATERIALS AND METHODS

### Single cell analysis of PDAC tissues and TCGA datasets

We analyzed a publicly available single-cell RNA-Seq dataset from human PDAC tissues (32). Data processing and analysis were performed using Bioturing’s Talk2Data V4 platform. To explore gene expression in specific cell populations, we utilized the sub-clustering function.

### Siglec-10 staining on immune cells

Monocytes isolated from healthy donors were differentiated into either immune-activating or immunosuppressive macrophages. For immune-activating conditions, monocytes were cultured with M-CSF (50 ng/mL) for 6 days, followed by stimulation with LPS (10 ng/mL) and IFN-γ (20 ng/mL) along with M-CSF (50 ng/mL) for 24 hours on day 6. For immunosuppressive differentiation, monocytes were treated with M-CSF (50 ng/mL) for 3 days, then stimulated with M-CSF (50 ng/mL) and IL-4 (20 ng/mL) from day 3 to day 8. Siglec-10 expression was assessed on monocytes at day 0 and on macrophages at their respective time-points. Cells were detached using TrypLE, stained with APC-conjugated anti-Siglec-10 antibody for 30 minutes, and washed twice with PBS. Samples were analyzed on a BD FACS Symphony A3 flow cytometer. Data analysis was performed using FlowJo (RRID:SCR_008520) version 9.

### Siglec-10 ligand and CD24 staining on cancer cells

Siglec-10 ligand expression was detected using either a recombinant chimeric protein consisting of the Siglec-10 binding region fused to human IgG Fc domains (Siglec-10-Fc, R&D Systems) or a biotin-tagged Siglec-10 mutant (R119A) (Kactus, catalog# SIG-HM411). To form the Siglec-10-Fc/anti-human IgG-BV421 complex, Siglec-10-Fc (2.5 μg/mL) and BV421-conjugated anti-human IgG were incubated on ice for 1 hour. Cancer cells were detached using TrypLE Express, washed, pelleted, and adjusted to a density of 1 × 10 cells/mL. The cells were then resuspended in the Siglec-10-Fc/anti-human IgG-BV421 precomplex solution together with CD24-Alexa 488 antibody. To form the Siglec-10 mutant (R119A)-streptavidin conjugate, biotin-tagged Siglec-10 mutant (R119A) (2.5 µg/test) was incubated with APC-streptavidin on ice for 1 hour. Cancer cells were detached using TrypLE Express, washed, pelleted, and adjusted to a density of 5 × 10 cells/test. The cells were then resuspended in the Siglec-10 mutant (R119A)/streptavidin-APC precomplex solution together with CD24-Alexa 488 antibody. After a 30-minute incubation at 4 °C, the cells were pelleted by centrifugation at 300 × g for 5 minutes and washed twice. Samples were analyzed on a BD FACS Symphony A3 cytometer, acquiring at least 30,000 events per sample. Data analysis was performed using FlowJo v9.

### Pull-Down and mass spectrometry analysis of Siglec-10 ligands on PDAC cells

To identify Siglec-10 ligands, we employed a proximity labeling approach based on the tyramide radicalization principle (33). Briefly, recombinant Siglec-10-Fc fusion protein (10 μg) was incubated with an anti-human Fc HRP conjugate (5 μg), a monoclonal antibody against human Fc tag directly conjugated to horseradish peroxidase (HRP), on ice for 60 minutes to facilitate complex formation. PDAC cell lines (AsPC-1 (RRID:CVCL_0152), BxPC-3 (RRID:CVCL_0186), MiaPaCa-2 (RRID:CVCL_0428), PANC-1 (RRID:CVCL_0480)) were incubated with the Siglec-10-HRP complex on ice for 1 hour (20 × 10 cells per reaction). After incubation, cells were washed twice with 10 mL of 140 mM NaCl in 20 mM Tris-HCl buffer (pH 8.0, TBS). The cells were then treated with 10 μM biotin tyramide and 10 mM H O in TBS at room temperature for 10 minutes. Following the reaction, cells were washed three times with TBS and lysed in 200 μL of lysis buffer (50 mM Tris-HCl, pH 8.0, 150 mM NaCl, 1% Nonidet P-40, 0.5% sodium deoxycholate, 0.1% SDS) supplemented with a protease inhibitor cocktail. Lysis was performed on ice for 30 minutes. A Siglec-5 control and a secondary antibody-only control were included for each cell line. The lysates were cleared by centrifugation at 15,000 × g for 30 minutes and used for the purification of biotinylated proteins. Biotinylated proteins were purified using streptavidin-functionalized paramagnetic beads. One milligram of Dynabeads Streptavidin-MyOne C1 (Thermo Fisher Scientific) was mixed with 100 μL of cleared lysate and incubated at room temperature for 1 hour with constant shaking. The beads were collected using a magnet (DynaMag-2, Thermo Fisher Scientific) and washed extensively with D-PBS containing 0.1% SDS. The samples were reduced with tris(2-carboxyethyl)phosphine, alkylated with iodoacetamide, and digested on-bead with trypsin (Promega). Tryptic digests were cleaned using BioPureSPN C18 spin columns (Nest Group) and analyzed by liquid chromatography-tandem mass spectrometry (LC-MS/MS) using a 1.5-hour LC gradient on a Thermo Q Exactive Plus mass spectrometer (34). MS data were searched with full tryptic specificity against the UniProt human proteome (downloaded on 8/21/2023) and a contaminant database using MaxQuant 2.4.7.0 (35) (RRID:SCR_014485). Protein and peptide false discovery rates were set at 1%. A total of 4,044 proteins were identified, which were further filtered based on a two-fold enrichment of Siglec-10 compared to the Siglec-5 control.

### Surface Plasmon Resonance (SPR)

SPR experiments were performed to evaluate the binding of various recombinant proteins to Siglec-10. Recombinant Siglec-10 Fc protein was immobilized on a protein A/G sensor chip at three different surface densities (∼200 response units [RU], ∼600 RU, and ∼1000 RU). One flow cell was left blank to serve as a reference control for non-specific binding to the chip surface. The following recombinant proteins were tested: CD47 (His tag, Acrobiosystems, Catalog# CD7-H5227), CD59 (His, Avitag, Acrobiosystems, Catalog# CD9-H82E3), NT5E/CD73 (His, Avitag, Acrobiosystems, Catalog# CD3-H82E3), ITGB6 (C-Myc/DDK, Origene, Catalog# TP317387), ITGA3 (C-Myc/DDK, Origene, Catalog# TP320975), and ITGB1 (C-Myc/DDK, Origene, Catalog# TP303818). All test proteins were desalted into running buffer before analysis. Proteins were tested at two concentrations (100 nM and 1000 nM), except for ITGA3, which was tested at 30 nM and 300 nM. For each injection, the association phase was set to 120 seconds and the dissociation phase to 600 seconds, with a flow rate of 25 µL/min. After each binding cycle, the Siglec-10/protein complex was dissociated from the chip surface using 20 mM glycine (pH 2.0), and fresh Siglec-10 protein was re-immobilized to maintain consistent binding capacity across injections.

### Sialidase treatment and lectin staining

ITGA3 and ITGB1 proteins were treated with Sialidase A (AdvanceBio Sialidase A; Agilent; catalog# GK80040) following the manufacturer’s instructions. Briefly, 5 μg of ITGA3 and ITGB1 glycoproteins were combined with 14 μL of deionized water, 4 μL of 5× Reaction Buffer, and 2 μL of Sialidase A. The reaction mixture was incubated at 37°C for 1 hour. To verify the removal of sialic acid, a lectin microarray platform was used to profile sialic acid-dependent glycan structures. This array utilizes a panel of immobilized lectins, each with known glycan structure binding specificity. Both sialidase-digested and undigested proteins were labeled with Cy3 dye (Sigma-Aldrich) and hybridized to the lectin microarray. The lectin chips were scanned for fluorescence intensity at each lectin-coated spot using an evanescent-field fluorescence scanner (Rexxam Co., Ltd.). All samples were run in triplicate, and the average fluorescence intensity of the triplicates was used for analysis. Data were normalized using the global normalization method.

### CRISPR/Cas9-mediated knockout of ITGA3 in PDAC Cells

Knockout of ITGA3 was performed using a synthetic guide RNA (gRNA) targeting ITGA3 (Horizon, Catalog# CM-004571-01; Target Sequence: CCAGGGTACACGATGCAGGT). A non-targeting gRNA served as a control (catalog# U-009502-01-02). CRISPR ribonucleoprotein (crRNP) complexes were assembled; 1 µL gRNA was combined with 1 µL tracrRNA and incubated at 37°C for 30 minutes to allow duplex formation. 1.6 µL of Poly-L-glutamic acid sodium salt (100mg/mL) was added to the duplex solution followed by the slow addition of recombinant Cas9 protein (2 µL). The final crRNP complex was incubated at 37°C for an additional 15 minutes to complete assembly. Assembled crRNPs (5.6 µL per reaction) were aliquoted into PCR strip tubes and stored on ice prior to transfection. PANC-1 cells were prepared at a density of 2 × 10 cells per reaction. Electroporation was performed using the Lonza 4D-Nucleofector system with the SE Cell Line 4D-Nucleofector X Kit (Lonza), following the manufacturer’s protocol. A 20 µL nucleofection buffer was prepared by combining 3.6 µL of SE Supplement with 16.4 µL of SE Solution. Cells were suspended in nucleofection buffer and mixed with 3.5 µL of the crRNP complex prior to nucleofection using program DN-100. Immediately following electroporation, 100 µL of pre-warmed complete growth medium was added to each reaction for recovery. Cells were then transferred to 6-well plates containing 2 mL of medium per well and incubated under standard conditions for 48 hours. Knockout efficiency was assessed 72 hours post-transfection by flow cytometry. Cells were stained with an APC-conjugated anti-ITGA3 antibody (Biolegend; catalog# 343808) and analyzed for surface expression. A reduction in ITGA3 signal compared to non-targeting control indicated successful gene knockout. Edited cell populations were maintained in culture and routinely monitored by flow cytometry at regular intervals to confirm stable loss of ITGA3 expression.

### Macrophage differentiation and in vitro phagocytosis assay

Macrophages were differentiated as described previously (20,36). Briefly, monocytes from healthy donors were differentiated into macrophages by culturing them for 7–9 days in Iscove’s Modified Dulbecco’s Medium (IMDM) supplemented with 10% AB human serum (Life Technologies). During the initial 3-4 days, macrophages were stimulated with 50 ng/mL M-CSF. Subsequently, M-CSF (50 ng/ml) and IL-4 (20 ng/mL) were added to support differentiation, and this stimulation was maintained until the macrophages were used on Days 7-9. PDAC cells were treated with TrypLE Express and labeled with pHrodo Red, SE (Thermo Fisher Scientific) according to the manufacturer’s protocol. Labeling was performed at a concentration of 250 ng/mL in PBS for 10 minutes at 37°C, followed by two washes with 1X PBS. Healthy donor-derived macrophages were harvested using TrypLE Express, and 25,000 macrophages were seeded into black 96-well flat-bottom plates, allowing them to adhere for 30 minutes at 37°C. After adhesion, 25,000 pHrodo Red-labeled PDAC cells were added in serum-free IMDM. The plates were incubated at 37°C, and phagocytosis was monitored by imaging every hour using an Incucyte (Sartorius). The initial image (t = 0) was captured within 30 minutes of co-culture. Phagocytosis was quantified by measuring the total red area (µm² per well) across technical replicates for each donor. Thresholds for identifying pHrodo Red-positive events were determined based on intensity measurements from labeled cells in the absence of macrophages.

### ITGA3^low^ and ITGB1^low^ PDAC cell enrichment

PDAC cells were incubated with anti-human ITGA3 and ITGB1 antibodies (Clone ASC-1 and TS2/16, respectively, from Thermo Fisher Scientific) at a concentration of 1 μg/mL in 1× PBS at 4°C for 15 minutes. The cells were then washed twice and incubated with 20 μL of anti-mouse MicroBeads (Miltenyi Biotec, catalog# 130-048-402) for 15 minutes at 4°C. Following another wash, the cells were loaded onto pre-equilibrated LS columns (Miltenyi Biotec, catalog# 130-042-401) in accordance with the manufacturer’s protocol. After washing, the cells in the eluate fraction were pelleted, resuspended in serum free IMDM, and subsequently used for in vitro cell-killing assays.

### Siglec-10 monoclonal antibody production

Monoclonal antibodies targeting the extracellular domain (ECD) of Siglec-10 were developed and characterized. Ten BALB/c (RRID:MGI:2683685) and C57BL/6 (RRID:IMSR_JAX:000664) mice were initially immunized with 50μg of recombinant Siglec-10 protein (GenScript), followed by four booster immunizations of 25μg each at two-week intervals. Blood samples were collected after each immunization, and serum antibody titers were assessed using indirect ELISA. For ELISA, 96-well plates were coated with Siglec-10 antigen or Siglec-5 antigen (1μg/mL in PBS) and incubated overnight at 4°C. Plates were blocked with 1% BSA in PBS to prevent nonspecific binding. Diluted serum samples were incubated for 1 hour at 37°C, followed by washes and incubation with horseradish peroxidase (HRP)-conjugated goat anti-mouse IgG secondary antibody. The reaction was developed using tetramethylbenzidine (TMB) substrate, stopped with acid, and the absorbance was measured at 450 nm. Mice with serum OD450 values exceeding 1.0 at dilutions ≥1:8,000 were selected for hybridoma generation. Splenocytes from selected mice were fused with SP2/0 myeloma cells via electrofusion. The hybridomas were cultured in HAT medium, and supernatants were screened for Siglec-10-specific antibodies using same ELISA protocol described above. Hybridoma clones were also tested for specificity by flow cytometry (FACS) using CHO-K1 cells (RRID:CVCL_0214) expressing Siglec-10 or Siglec-5 (as a control). Positive hybridoma supernatants demonstrated significant fluorescence with Siglec-10-expressing cells compared to controls. For FACS, 50 μL of a CHO-K1 cell suspension (1 × 10 cells per well) was incubated with hybridoma supernatants at 4°C for 30 minutes. After washing, Alexa Fluor 488-conjugated goat anti-mouse IgG secondary antibody (Jackson ImmunoResearch) was added, and samples were analyzed using a flow cytometer. Clones with high median fluorescence intensity (MFI) against Siglec-10 cells were selected for subcloning.

Selected hybridoma clones underwent limiting dilution for subcloning. Subclones were cultured, and supernatants were collected for antibody purification. Antibodies were purified using Protein A chromatography and assessed for purity by SDS-PAGE. Concentrations were determined using UV spectrophotometry. The specificity of purified antibodies was validated using the same ELISA and FACS protocols described above against both Siglec-10 and Siglec-5 (as a control). Antibodies from positive clones were sequenced. RNA was extracted from hybridoma cells using the RNeasy Isolation Kit, and antibody sequences were determined via RT-PCR using universal primers. Sequenced antibodies were expressed recombinantly in CHO-S cells and purified. Functional characterization of these recombinant antibodies was performed using the same ELISA and FACS assays described above.

### Co-culture assays with monocyte-derived macrophages and CD8^⁺^ T cells

Human CD8^⁺^ T cells and monocytes were obtained from the Human Immunology Core at the University of Pennsylvania. CD8^⁺^ T cells were labeled with 5 μM CellTrace Violet (Thermo Fisher Scientific) and co-cultured with autologous monocytes at a 1:2 ratio (T cell:monocyte) for 5 days in the presence of anti-CD3/CD28 Dynabeads (Thermo Fisher Scientific) for polyclonal T cell activation. At the end of the culture period, cells were harvested and analyzed using a BD FACSymphony flow cytometer (BD Biosciences). Proliferation was assessed based on CellTrace dye dilution. For GranzymeB staining, cells were fixed and permeabilized using True nuclear transcription factor buffer set (Biolegend, Catalog# 424401), according to the manufacturer’s instructions. Flow cytometry data were analyzed using FlowJo software (v10.6.1, TreeStar), and statistical analyses and graphical representations were performed using GraphPad Prism (v10.0.0).

### Cancer associated fibroblasts (CAFs), macropahges, and cancer cell triple co-culture assays

CAFs were cultured in DMEM supplemented with 10% FBS. Macrophages were differentiated from healthy donor monocytes as previously described. After differentiation, macropahges were detached using TrypLE Express, and 25,000 cells per well were seeded into a 96-well plate in the presence of either a Siglec-10 blocking antibody or an isotype control (10 µg/well). TAMs were allowed to adhere for 1 hour at 37°C. CAFs and PANC-1 cells were detached using TrypLE Express. PDAC cells were labeled with pHrodo Red as previously mentioned. Once TAMs adhered, 25,000 pHrodo Red-labeled PDAC cells and 10,000 CAFs were added per well in serum-free IMDM. Plates were incubated at 37°C, and phagocytosis was monitored hourly using an Incucyte live-cell imaging system (Sartorius).

### Experiments using xenograft mouse model

NSG (NOD.Cg-Prkdc^scid^ Il2rg^tm1Wjl^/SzJ) mice (RRID:BCBC_4611) were anesthetized with isoflurane, and the hair on the right flank was shaved. A 100 µL suspension of AsPC-1 cells (2 × 10 cells per mouse) was prepared by mixing the cells with Matrigel and 1× PBS at a 1:1 ratio, which was then injected subcutaneously into each mouse. After one-week, subcutaneous tumors of approximately 40 mm³ had developed. The mice were randomly assigned to two groups: isotype control or Siglec-10 antibody treatment. Monocyte-derived macrophages were incubated with 200 µg of either isotype or Siglec-10 antibody for 30 minutes. Each mouse received intravenous injections of antibody-incubated macrophages three times per week for three weeks. Tumor size was measured using a vernier caliper, and tumor volume was calculated using the formula: tumor volume = ½ (length × width²). All mice were maintained inappropriate environmental conditions. All animal protocols were approved by Wistar Institutional Animal Care and Use Committee (protocol # 201363). Euthanasia was performed using carbon dioxide (CO2) delivered as bottled gas, which is listed among the acceptable agents for euthanasia of laboratory rodents in the American Veterinary Medical Association (AVMA) Guidelines for the Euthanasia of Animals: 2013 Edition.

### RNA-Seq analysis of macrophages

Subcutaneous tumors were excised from each of the xenograft mice, and human cells were sorted by FACS. Total RNA was extracted from the sorted human macrophages using the Single Cell RNA Purification Kit (Norgen). RNA quality was assessed using the TapeStation High Sensitivity RNA ScreenTape (Agilent). Libraries were prepared using the Low Input RNA Library Prep Kit (Takara Biosciences) with 500 ng of DNase I-treated total RNA. Final library quality control was performed using the Bioanalyzer High Sensitivity DNA Kit (Agilent). Next-generation sequencing (NGS) was conducted with 100 bp paired-end reads on a NovaSeq 6000 system (Illumina) using the SP v1.5 200-cycle kit. RNA-seq analysis was performed by Bencos Research Solutions Pvt. Ltd. (Mumbai) using their proprietary TWINE BI platform. For the analysis, the RNA-seq data were aligned to the hg19 human genome using STAR-aligner (v 2.7.10a), and salmon (v 1.10.1) was used to estimate read counts and RPKM values. Raw counts were analyzed for differential expression using DESeq2 (RRID:SCR_000154).

### Experiments using human Siglec-10 transgenic mouse model

B-hSIGLEC10 mice [C57BL/6N-(Siglecgtm2(SIGLEC10) Bcgen/Bcgen)] were obtained from Biocytogen (Catalog # 110843). In this genetically engineered strain, exons 3–10 of the murine *Siglec-G* gene, encoding the extracellular domain, were replaced with the corresponding human *SIGLEC10* exons, resulting in a humanized extracellular domain. Knockout of mouse Siglec-G and knock-in expression of human SIGLEC10 were confirmed by flow cytometry analysis using the following fluorophore-conjugated antibodies: F4/80-BUV805 (BM8, ThermoFisher, 368-4801-80), CD3-BV510 (OKT3, BioLegend, 317331), MHC-II-BV711 (M5/114.15.2, BioLegend, 107643), Ly-6C-Pacific Blue (HK1.4, BioLegend, 128013), CD11c-PE-Cy7 (N418, BioLegend, 117317), CD19-PE-Cy5 (6D5, BioLegend, 115509), mouse Siglec-G-PE (SH1, BioLegend, 163304), CD45-APC-Cy7 (30-F11, BioLegend, 103115), CD11b-Alexa Fluor 700 (M1/70, BioLegend, 101222), human Siglec-10-APC (5G6, BioLegend, 347606), and the LIVE/DEAD Fixable Blue Dead Cell Stain (ThermoFisher, L23105). B-hSIGLEC10 mice were anesthetized using isoflurane, and the hair on the right flank was removed. A 100 µL cell suspension containing 3 × 10 KPCY (2838c3) tumor cells was prepared in a 1:1 mixture of Matrigel and 1× PBS. The suspension was injected subcutaneously into the right flank of each mouse. The cell line was tested for contamination by Center for Comparative Medicine at Northwestern University. After 5 days, tumors reached an approximate volume of 40 mm³, at which point mice were randomized into two treatment groups: isotype control or anti-Siglec-10 antibody. Mice received daily intraperitoneal injections of either Siglec-10 blocking antibody or respective isotype until the study endpoint. Tumor growth was monitored using vernier calipers, and tumor volume was calculated using the formula: tumor volume = ½ (length × width²).

### Data availability

All data are available within the manuscript and its figures and/or available upon request.

## RESULTS

### Single-cell analysis of the PDAC TME showed that Siglec-10 is predominantly expressed in multiple myeloid cell subtypes

We began our investigation by asking which PDAC TME cells expressed Siglec-10. We analyzed a publicly available single-cell RNA sequencing (scRNAseq) dataset from human PDAC tumors (**Fig. 1a**) (32) and found that Siglec-10 is expressed in several myeloid cell subsets within the PDAC TME. Specifically, 60% of Siglec-10^⁺^ cells were macrophages, followed by myeloid-derived suppressor cells (MDSCs) and dendritic cells (DCs) (**Fig. 1b**). Turning to which Siglecs were most highly expressed on these myeloid cells, we found that Siglec-10 had higher expression than the other Siglecs previously implicated in PDAC immune evasion, in particular Siglec-7, −9, and −15 (**Fig. 1c**).

**Figure 1.**
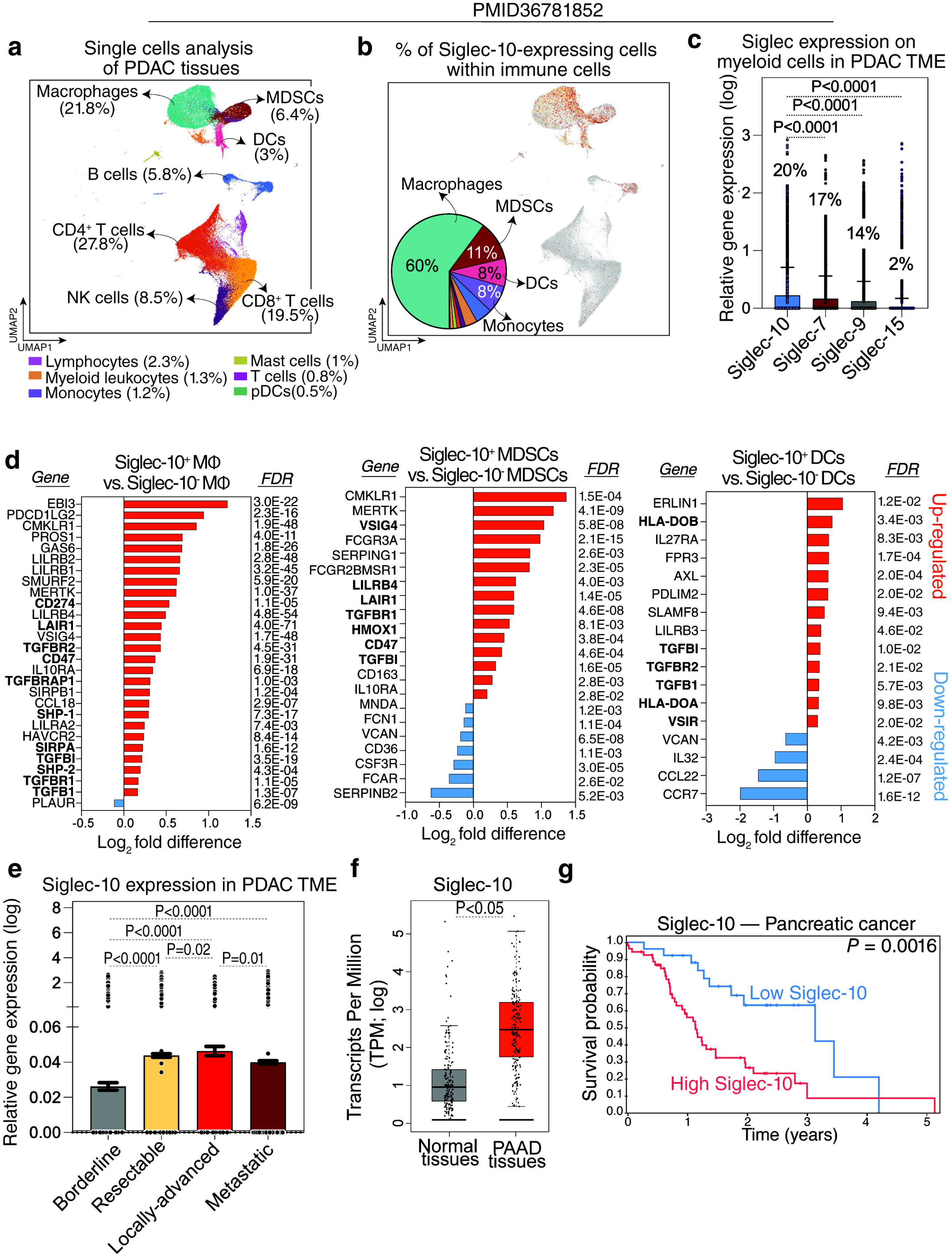
Siglec-10 is predominantly expressed on myeloid cells within the PDAC TME and is associated with worse disease progression. (**a**) UMAP visualization of immune cell populations within the PDAC TME. (**b**) UMAP of Siglec-10 expression and a pie chart illustrating the percentage of Siglec-10 cells within each myeloid cell subset. (**c**) A comparison of the expression levels of various Siglecs in myeloid cells within the PDAC TME. Kruskal– Wallis test with Dunn’s multiple comparisons correction. Means with standard deviations (SD) are shown. Percentages shown represent the proportion of myeloid cells expressing each of these Siglecs. (**d**) A comparison of the gene expression of the indicated genes between Siglec-10 and Siglec-10 macrophages, MDSCs, and DCs. Analysis was performed using the Bioturing’s Talk2Data V4 platform. (**e**) Siglec-10 expression in the PDAC TME across different disease states. Kruskal–Wallis test with Dunn’s multiple comparisons correction. Means with standard errors of the mean (SEM) are shown. (**f**) Comparison of Siglec-10 expression between normal tissues and pancreatic adenocarcinoma (PAAD) tissues from the TCGA dataset. Unpaired t test. (**g**) Survival analysis of pancreatic tumor patients in the Human Protein Atlas dataset showing the correlation between Siglec-10 expression and overall survival.

These findings suggest that Siglec-10 expression on TAMs, MDSCs, and DCs may contribute to their immunosuppressive phenotypes. Supporting this hypothesis, Siglec-10 cells exhibited elevated expression of genes linked to immunosuppressive pathways, particularly TGF-β signaling (22,37–42), and lower expression of pro-inflammatory and activation-associated genes. Siglec-10 macrophages also showed increased expression of SHP-1 and SHP-2, key mediators of Siglec-inhibitory signaling. In DCs, Siglec-10 cells exhibited upregulation of genes impairing antigen presentation, such as HLA-DOB and HLA-DOA (**Fig. 1d**). Additionally, Siglec-10 expression within the PDAC TME was associated with more advanced and metastatic disease (**Fig. 1e**).

To assess the clinical relevance of Siglec-10 expression in pancreatic cancer, we analyzed The Cancer Genome Atlas (TCGA; version 1, www.gepia.cancer-pku.cn) data for pancreatic adenocarcinoma (PAAD). PAAD tissues exhibited higher Siglec-10 expression than normal pancreatic tissues (**Fig. 1f**), and elevated Siglec-10 levels correlated with poorer survival outcomes in the Human Protein Atlas dataset (www.proteinatlas.org; **Fig. 1g**). In addition to examining Siglec-10 expression on macrophages, we investigated the potential role of its ligands, sialic acids, in PDAC progression. Sialic acids are added to glycoproteins through the activity of various glycan-related enzymes, including sialyltransferases (**Supplementary Fig. 1a**). We found that multiple genes involved in the synthesis of sialic acid ligands for Siglec-10 were significantly upregulated in PAAD tissues compared to normal tissues (**Supplementary Fig. 1b–c**), suggesting that Siglec-10–sialic acid interactions may contribute to PDAC progression.

Together, these data show that Siglec-10, a glyco-immune checkpoint, is highly expressed in the PDAC TME and is associated with disease progression and poorer prognosis. To validate these findings at the protein level, we differentiated primary human monocytes into macrophages using M-CSF, followed by IL-4 or LPS plus IFN-γ to generate immunosuppressive or immunoactivating phenotypes, respectively. Siglec-10 protein levels, measured by flow cytometry, were higher on immunosuppressive macrophages than on immunoactivating macrophages (**Supplementary Fig. 2**). Overall, these results suggest that Siglec-10 expression on myeloid cells within the PDAC TME is associated with an immunosuppressive phenotype.

### PDAC cells bind to Siglec-10 through sialic acid expressed on proteins other than CD24

Our next goal was to identify the glycoprotein ligand(s) of Siglec-10 on PDAC cells. CD24 was recently identified as a major glycoprotein ligand of Siglec-10 on breast and ovarian cancer cells (20). To test whether PDAC cells express CD24 and whether this expression coincides with Siglec-10 binding, we used a Siglec-10–Fc chimera protein followed by a secondary antibody, along with anti-CD24 antibodies, to detect Siglec-10 ligands and CD24 expression by flow cytometry on multiple PDAC cell lines as well as ovarian and breast cancer cells. Multiple human PDAC cell lines bound Siglec-10, indicating the presence of compatible sialylated glycoproteins on their surface (**Fig. 2a–f**). However, only a subset of these cells expressed CD24, with a large population of CD24-negative cells capable of binding Siglec-10 (**Fig. 2a–f**). This was in contrast to ovarian and breast cancer cells, which expressed high levels of CD24 that coincided with the presence of Siglec-10 ligands (**Fig. 2i–l**). Importantly, using a Siglec-10 protein containing the inhibitory R119A mutation, which abrogates sialic acid binding, we confirmed that Siglec-10 binding to PDAC cells is sialic acid–dependent (**Fig. 2**). These data suggest that PDAC cells engage Siglec-10 on myeloid cells through glycoproteins other than CD24.

**Figure 2.**
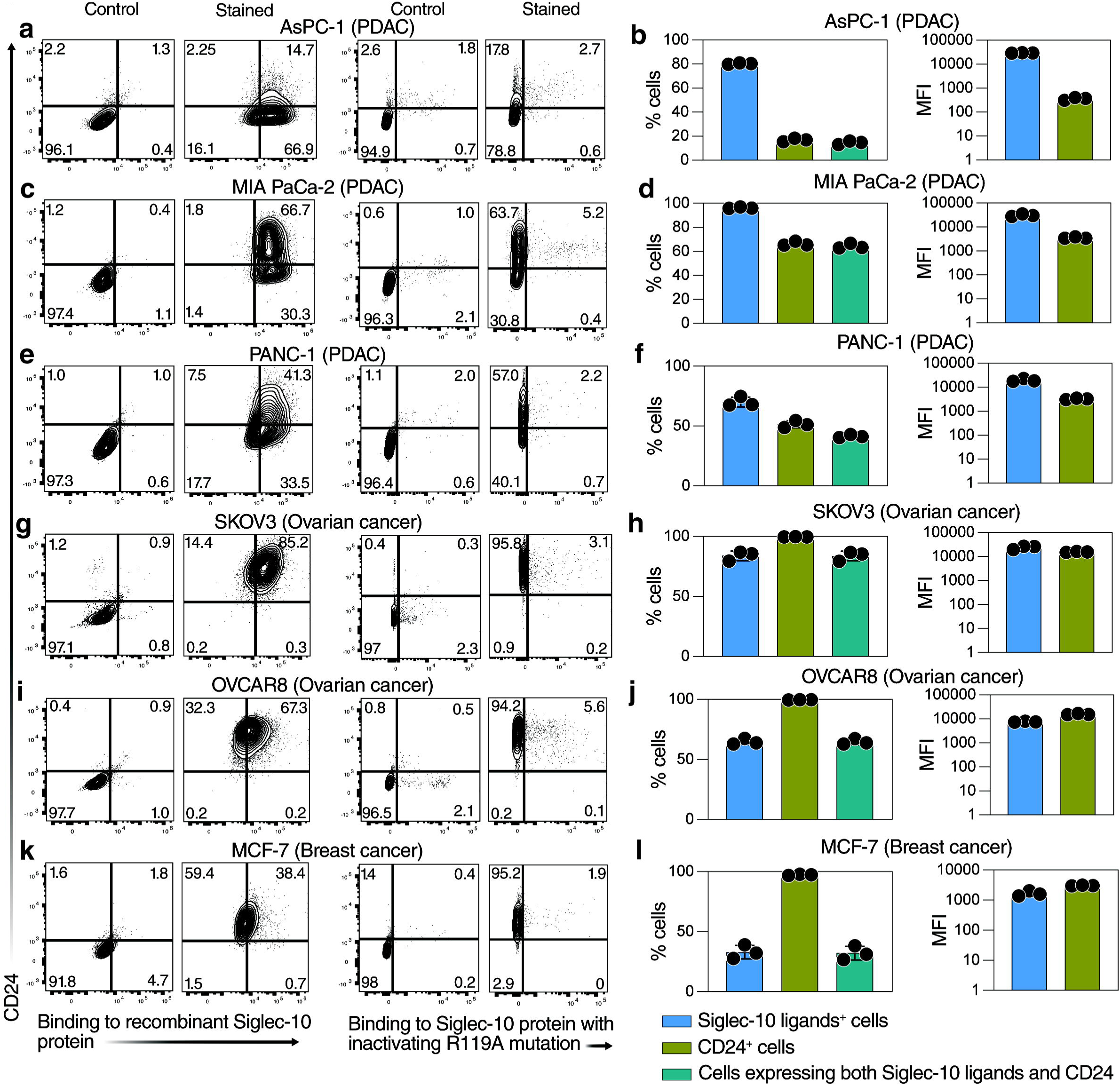
PDAC cell lines express Siglec-10 ligands on glycoproteins other than CD24. Flow cytometric analysis of Siglec-10 ligand and CD24 expression on (**a–f**) PDAC cell lines, (**g–j**) ovarian cancer cell lines, and (**k–l**) a breast cancer cell line. Siglec-10 ligands were detected using either (1) a recombinant Siglec-10-Fc chimeric protein and a secondary antibody (x-axis), with CD24 detected using a specific antibody (y-axis), or (2) a recombinant Siglec-10-Fc chimeric protein containing the inhibitory R119A mutation, which abrogates sialic acid binding (x-axis), with CD24 detected using a specific antibody (y-axis). Panels on the left (**a, c, e, g, i, k**) show representative flow plots. Panels on the right (**b, d, f, h, j, l**) show the percentage of positive cells and the median fluorescence intensity (MFI) for Siglec-10 ligands (using the non-mutated recombinant Siglec-10 protein), CD24, or cells expressing both. Means with SD are shown.

### Siglec-10 binds to ITGA3 and ITGB1, the subunits of α3β1 Integrin, on PDAC cells

To identify glycoproteins on PDAC cells that bind to Siglec-10, we performed a pull-down assay using recombinant Siglec-10 protein and cell-membrane proteins isolated from several human PDAC cell lines. Proteins bound to Siglec-10 were identified via mass spectrometry analysis (**Fig. 3a**). From a total of 4,044 proteins whose binding was enriched compared to a no-protein control (a control with anti-Fc HRP secondary antibody only), 110 proteins showed enrichment compared to Siglec-5 protein (a closely related Siglec used as a control). Of these, six proteins, CD47, CD59, CD73, ITGB6, ITGA3, and ITGB1, were significantly overexpressed in PAAD tissues compared to normal tissues based on TCGA data (**Fig. 3b**).

**Figure 3.**
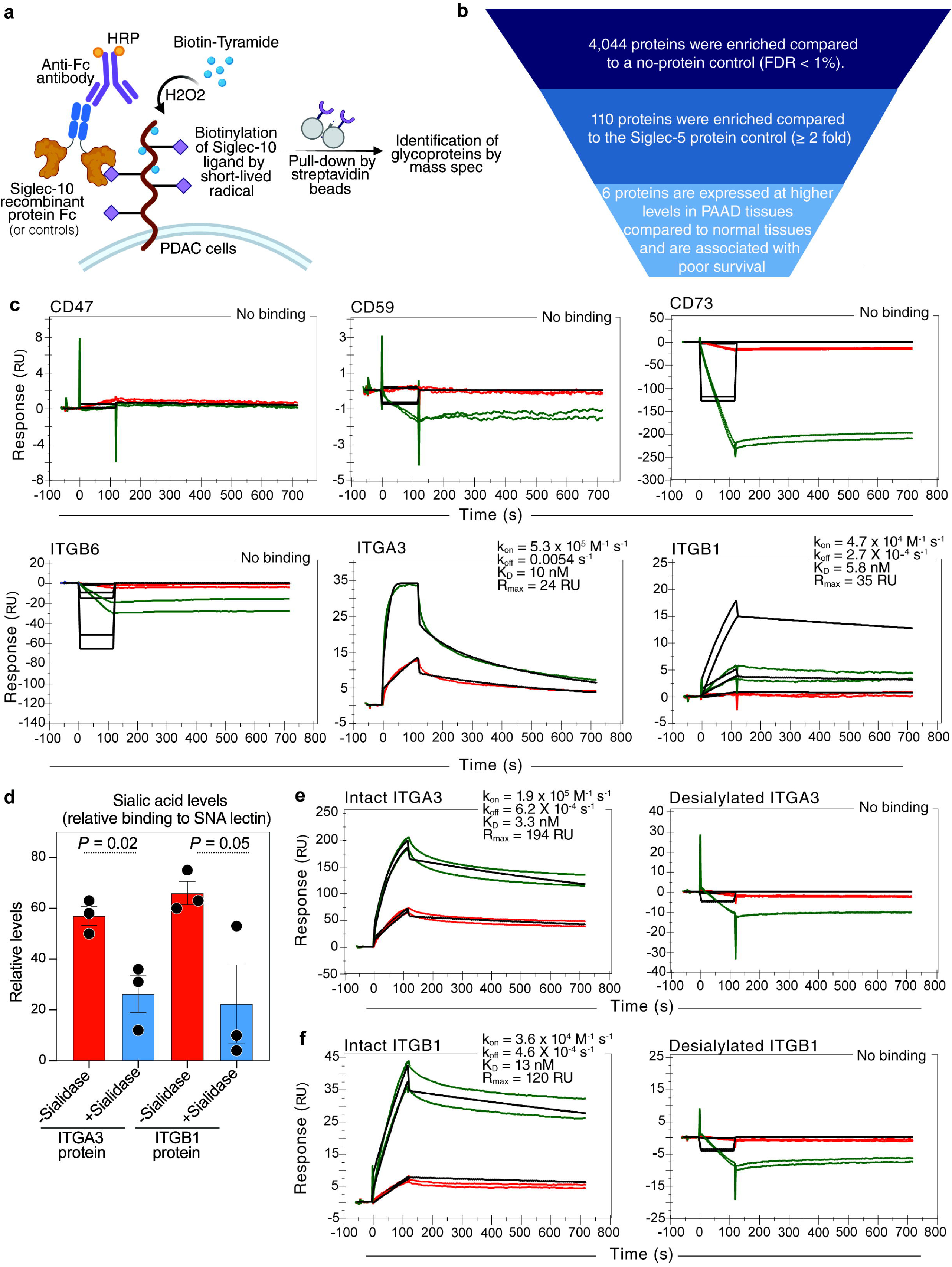
ITGA3 and ITGB1 are sialylated glycoprotein ligands of Siglec-10 glycoproteins on PDAC cells. **(a)** Experimental design for the pull-down of Siglec-10 ligands on PDAC cells. Recombinant Siglec-10 Fc (as well as a no-protein control or a Siglec-5 control) was allowed to bind its physiological ligands on the surface of PDAC cells, followed by an HRP-conjugated anti-Fc secondary antibody. In the presence of H□O□, HRP generated short-lived radicals that facilitated the transfer of biotin to proximal Siglec-10 ligands. Biotinylated Siglec-10 ligands were pulled down using streptavidin beads and identified by mass spectrometry. **(b)** A total of 4,044 proteins were identified, with enriched binding compared to a control using the anti-Fc antibody only without Siglec-10 protein. Of these, 110 proteins showed enrichment relative to a Siglec-5 control. Six proteins, CD47, CD59, CD73, ITGB6, ITGA3, and ITGB1, were significantly overexpressed in PAAD tissues compared to normal tissues in the TCGA dataset. **(c)** Response curves showing interactions between Siglec-10 and the six glycoproteins measured by surface plasmon resonance (SPR). Two concentrations (1000 nM, green; 100 nM, red) were tested for all glycoproteins, while ITGA3 was also tested at 300 nM (green) and 30 nM (red). **(d)** Binding of the SNA lectin (specific for sialic acid) to ITGA3 and ITGB1 recombinant glycoproteins was measured by a lectin array. Sialidase-treated glycoproteins (red bars) showed significantly reduced binding compared to untreated glycoproteins (blue bars). Unpaired t-tests. Means with SEM are shown. **(e)** SPR response curves comparing the binding of intact (sialylated) ITGA3 and desialylated ITGA3 to immobilized Siglec-10. **(f)** SPR response curves comparing the binding of intact (sialylated) ITGB1 and desialylated ITGB1 to immobilized Siglec-10.

To determine whether these six proteins bind directly to Siglec-10, we performed surface plasmon resonance (SPR) analysis with immobilized recombinant Siglec-10 protein using recombinant versions of these proteins (produced in human cell lines to maintain their glycosylation). Among the six candidates, only ITGA3 and ITGB1, the subunits of α3β1 integrin, exhibited strong binding affinity to Siglec-10 (**Fig. 3c**). To assess whether this binding is dependent on the sialic acid content of these glycoproteins, we treated ITGA3 and ITGB1 with sialidase to enzymatically remove sialic acid. The removal of sialic acid was confirmed using lectin staining with a sialic acid–specific lectin (SNA), which showed a significant reduction in sialic acid levels on ITGA3 and ITGB1 after sialidase treatment (**Fig. 3d**). Sialidase treatment abolished the ability of ITGA3 and ITGB1 to bind to Siglec-10 (**Fig. 3e-f**). These findings show that Siglec-10 binds to the sialic acid on ITGA3 and ITGB1.

### The expression of ITGA3 and ITGB1 on PDAC cells correlates with disease progression and prognosis

Having identified ITGA3 and ITGB1 as glycoprotein ligands that engage Siglec-10 to potentially suppress immune surveillance, we next investigated whether their expression is associated with PDAC progression and patient outcomes. First, we measured the surface expression of ITGA3 and ITGB1 on several human PDAC cell lines and observed consistently high levels across multiple lines (**Supplementary Fig. 3**). To evaluate their *in vivo* relevance, we analyzed TCGA data comparing gene expression in PAAD tissues to normal pancreatic tissues. While CD24 levels were similar between PAAD and normal tissues, ITGA3 and ITGB1 expression was significantly higher in PAAD samples (**Fig. 4a–c**). Furthermore, ITGA3 and ITGB1 expression positively correlated with elevated Siglec-10 levels (**Supplementary Fig. 4a,c**) and with higher expression of glycosylation-related genes such as ST3GAL1 and B3GNT3, which are responsible for adding sialic acids to glycoproteins (**Supplementary Fig. 4b,d**).

**Figure 4.**
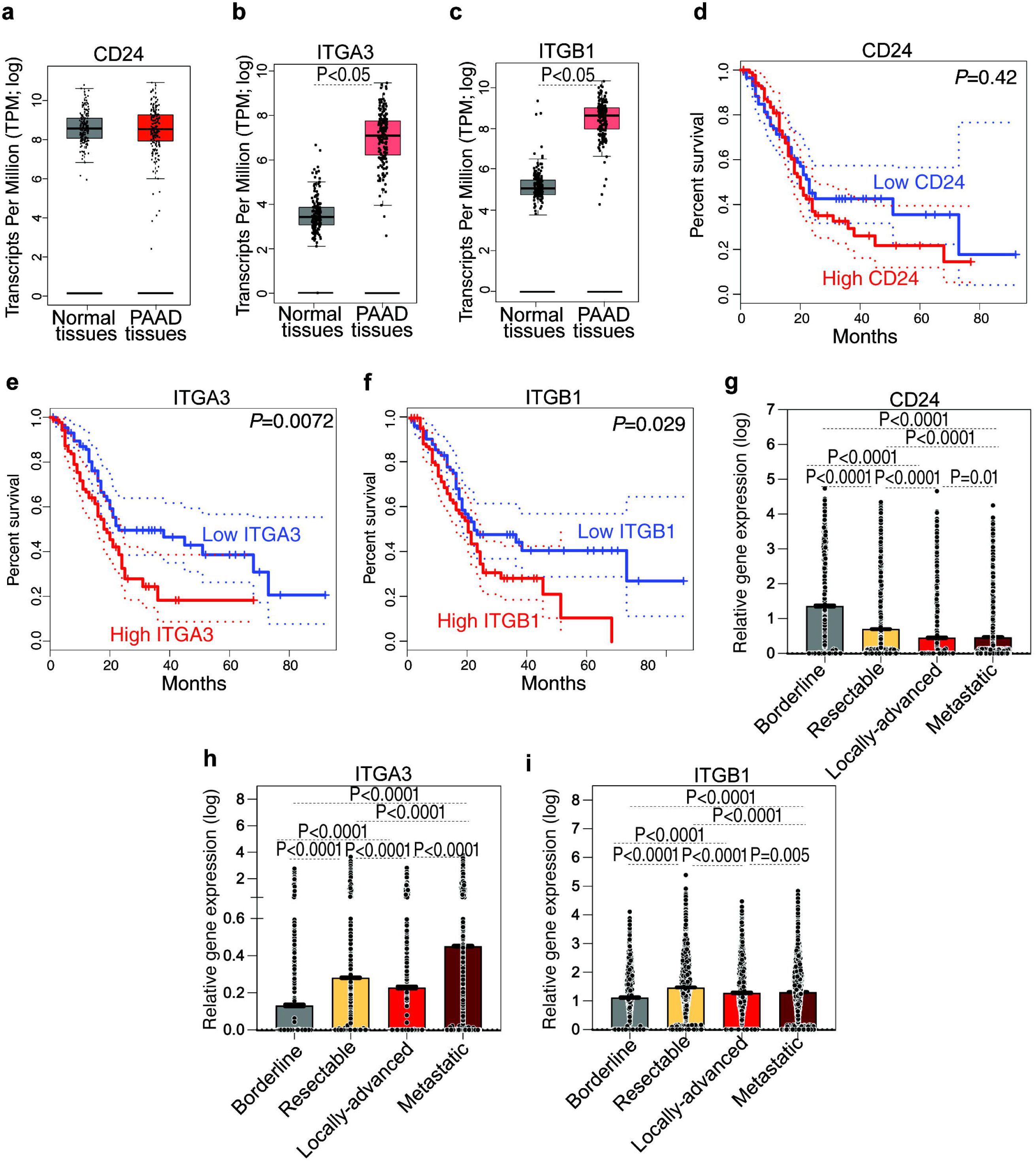
ITGA3 and ITGB1 are overexpressed in pancreatic cancer tissues and associated with poor survival. (**a–c**) Comparison of CD24 (a), ITGA3 (b), and ITGB1 (c) expression between normal tissues and PAAD tissues in the TCGA dataset. Unpaired t tests. (**d–f**) Survival analyses of pancreatic tumor patients in the TCGA dataset showing the correlation between CD24 (d), ITGA3 (e), and ITGB1 (f) expression and overall survival. (**g–i**) Expression of CD24 (g), ITGA3 (h), and ITGB1 (i) in the PDAC TME across different disease states. Kruskal–Wallis test with Dunn’s multiple comparisons correction. Means with SEM are shown.

Next, we examined the association of CD24, ITGA3, and ITGB1 expression with PDAC patient survival. While CD24 expression showed no correlation with survival outcomes (**Fig. 4d**), higher expression of ITGA3 (**Fig. 4e**) and ITGB1 (**Fig. 4f**) was significantly linked to worse prognosis. Finally, single-cell expression analyses within the PDAC TME revealed that elevated ITGA3 and ITGB1 expression correlated with more aggressive disease features, whereas CD24 expression did not (**Fig. 4g–i**). Together, these findings support the notion that ITGA3 and ITGB1 contribute to PDAC progression, potentially by engaging Siglec-10 and facilitating immune evasion.

### ITGA3 and ITGB1 facilitate PDAC cell evasion from macrophage-mediated phagocytosis

Given that (1) sialic acid on ITGA3 and ITGB1 in PDAC cells binds strongly to Siglec-10, (2) ITGA3 and ITGB1 expression is associated with PDAC progression and poorer prognosis, and (3) Siglec-10 interactions on macrophages inhibit their phagocytic activity, we next investigated whether PDAC cells with high ITGA3 or ITGB1 expression exhibit greater evasion of macrophage-mediated phagocytosis compared to cells with low expression of these glycoproteins.

We first employed the PDAC cell line PANC-1 and knocked out ITGA3 using CRISPR-Cas9. As shown in **Fig. 5a**, this effectively reduced cell-surface ITGA3 expression by over 95%, as measured by flow cytometry. We then examined whether this reduction enhanced the susceptibility of PDAC cells to macrophage-mediated phagocytosis. ITGA3-KO cells and non-targeting gRNA control cells were labeled with pHrodo dye and co-cultured with macrophages differentiated from monocytes of multiple healthy donors using M-CSF and IL-4 (to generate immunosuppressive monocyte-derived macrophages). We then assessed phagocytosis using live-imaging staining over time (**Fig. 5b**). As shown in **Fig. 5c**, ITGA3-KO PDAC cells were significantly more susceptible to macrophage-mediated phagocytosis than control PDAC cells with high ITGA3 expression.

**Figure 5.**
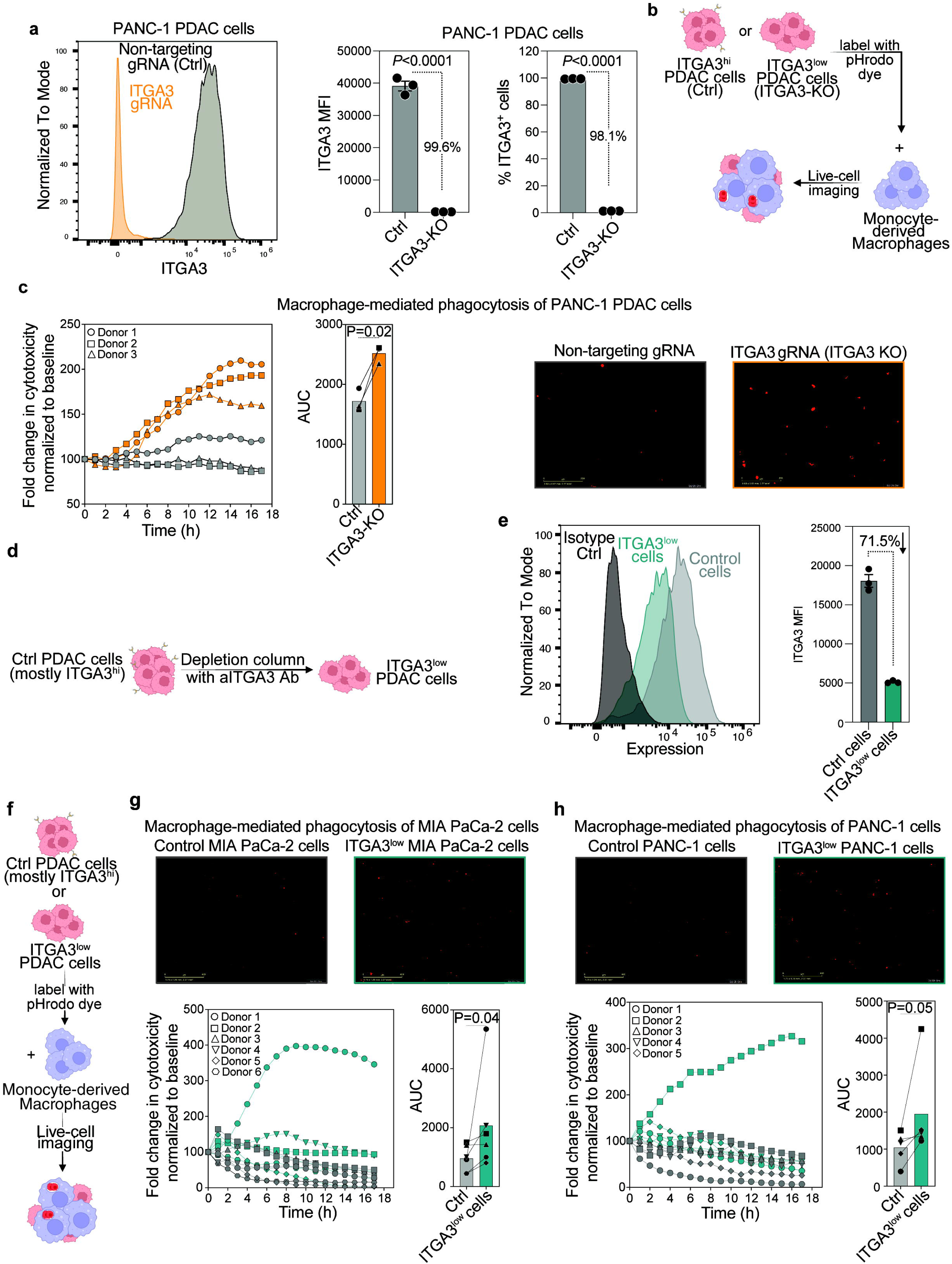
ITGA3-expressing PDAC cells evade macrophage-mediated phagocytosis. (**a**) ITGA3 knock-down in PANC-1 PDAC cells was achieved using CRISPR-Cas9. Flow cytometry confirmed the reduction of ITGA3 expression in knockout cells (ITGA3-KO) compared to non-targeting gRNA-treated cells and untreated controls. Data from three independent replicates show the percentage of ITGA3-expressing cells and the mean fluorescence intensity (MFI). Means and SEM are shown. Unpaired t-tests. (**b**) Experimental schematic illustrating co-culture of red-labeled PDAC cells (ITGA3-KO or non-targeting gRNA-treated) with monocyte-derived macrophages. Phagocytosis was monitored over time using live-cell imaging on the Incucyte Live-Cell Analysis System. (**c**) (left) Phagocytosis over time and area under the curve (AUC). Data show macrophage-mediated phagocytosis of PDAC cells, normalized to the 0-hour time point. Paired t-tests. N = 3 donors, with experiments performed in triplicate for each donor. (right) Representative images showing increased red fluorescence as an indicator of enhanced phagocytosis. (**d**) Schematic of ITGA3^low^ PDAC cell enrichment. ITGA3^high^ PDAC cells were separated using anti-ITGA3 antibody-coated columns, and the flow-through, containing ITGA3^low^ cells, was collected. (**e**) Flow cytometry analysis of ITGA3 expression in MIA PaCa-2 cells after enrichment for ITGA3^low^ cells and comparison with non-enriched controls (Ctrl cells). (**f**) Experimental schematic illustrating the phagocytic assay setup, where monocyte-derived macrophages were co-cultured with either ITGA3^high^ (control) or ITGA3^low^ PDAC cells. (**g-h**) Phagocytosis analysis of ITGA3^high^ (control) versus ITGA3^low^ MIA PaCa-2 (g) or PANC1 (h) cells by macrophages derived from different donors. N = 5–6, with experiments performed in triplicate for each donor. Top: Representative images showing increased red fluorescence as an indicator of enhanced phagocytosis. Bottom left: Live imaging data from individual donors (each symbol represents one donor’s data done in triplicate per donor). Bottom right: AUC data from multiple donors, showing significantly higher phagocytic activity with ITGA3^low^ cells. Paired t-tests.

To validate these results, we used the PDAC cell lines MIA PaCa-2 and PANC-1 and employed antibody-coated columns targeting ITGA3 to enrich for ITGA3^low^ cells (cells not bound by the columns) (**Fig. 5d**). Flow cytometry confirmed a significant enrichment of ITGA3^low^ cells compared to controls, which consisted predominantly of ITGA3^high^ cells that passed through the columns. **Fig. 5e** illustrates an example where depletion of ITGA3^high^ cells in MIA PaCa-2 resulted in over a 70% reduction in ITGA3 expression. Consistently and in subsequent phagocytosis assays (**Fig. 5f**), PDAC cells with high ITGA3 expression evaded macrophage-mediated phagocytosis more effectively than ITGA3^low^ cells (**Fig. 5g-h**). Similar results were observed when depleting ITGB1^high^ cells (**Supplementary Fig. 5**). Together, these findings indicate that ITGA3 and ITGB1 significantly contribute to the immune evasion of PDAC cells by inhibiting macrophage-mediated phagocytosis.

### Disrupting Siglec-10 interactions, via monoclonal antibodies, enhances macrophage-mediated phagocytosis of PDAC cell and prevent myeloid cell–mediated inhibition of T cell proliferation and activation in vitro

Our findings thus far suggest that Siglec-10 on macrophages binds to multiple glycoprotein ligands on PDAC cells, including ITGA3, ITGB1, and potentially others. This binding inhibits the phagocytic capacity of macrophages against PDAC cells (**Fig. 6a**). Therefore, directly targeting Siglec-10 itself, rather than its individual ligands, may be necessary to enhance macrophage-mediated phagocytosis of PDAC cells. Such an approach would disrupt interactions between Siglec-10 and sialic acids across its various glycoprotein ligands. However, the lack of potent, commercially available anti-Siglec-10 antibodies has necessitated the development of these antibodies in-house.

**Figure 6.**
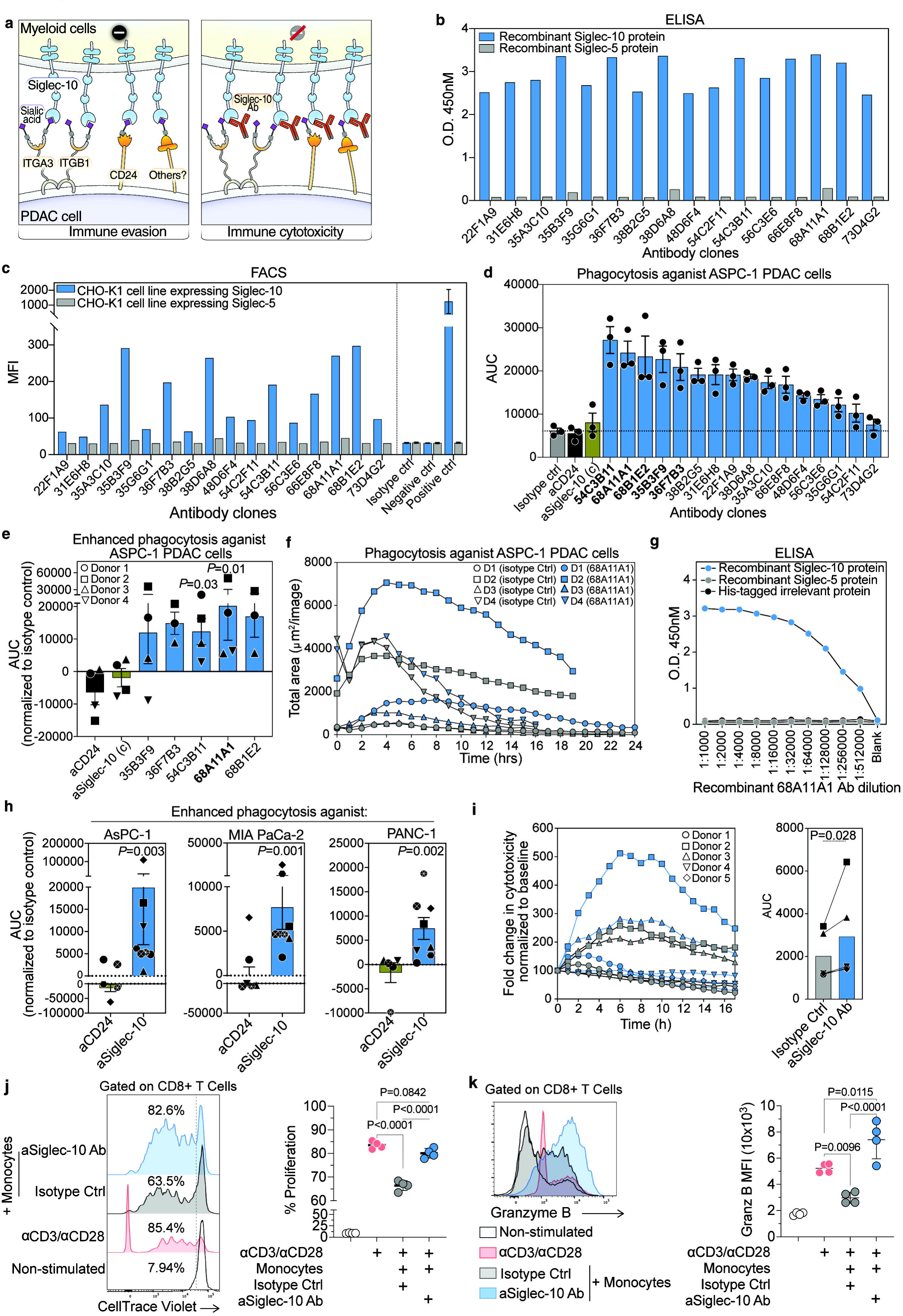
Development of Siglec-10 blocking antibodies to enhance macrophage-mediated phagocytosis of PDAC cells and prevent myeloid cell–mediated inhibition of T cell proliferation and activation *in vitro*. (**a**) Schematic model illustrating Siglec-10-mediated suppression of macrophage phagocytosis. In the left panel, Siglec-10 on macrophages binds to glycan ligands on PDAC cells, including ITGA3, ITGB1, and CD24, triggering inhibitory signaling and suppressing phagocytosis. In the right panel, blocking Siglec-10 with an antibody prevents inhibitory signaling and enhances macrophage phagocytic capacity. (**b**) ELISA screening of recombinant antibodies from the top hybridoma clones for Siglec-10 binding. Binding to immobilized Siglec-10 (blue) and Siglec-5 (gray) proteins is shown. (**c**) Flow cytometric analysis of antibody selectivity, showing binding to CHO-K1 cells expressing Siglec-10 (blue) but not Siglec-5 (gray). (**d**) AUC analysis of *in vitro* phagocytosis assays screening various Siglec-10 antibody clones, along with commercially available anti-CD24 and anti-Siglec-10 antibodies, for their ability to enhance macrophage-mediated phagocytosis of AsPC-1 PDAC cells. Means with SEM are shown. (**e**) AUC analysis of the *in vitro* phagocytosis assay using the top-performing Siglec-10 antibody clone with macrophages differentiated from monocytes of four healthy donors. Statistical significance was determined using Friedman’s ANOVA test. Means with SEM are shown. (**f**) Time-course analysis of the *in vitro* phagocytosis assay comparing the top Siglec-10 blocking antibody clone (68A11A1, blue) with the isotype control (gray). Data represent n = 4 independent experiments. (**g**) ELISA-based binding analysis of the recombinant 68A11A1 antibody to immobilized recombinant Siglec-10 and Siglec-5 proteins across different dilutions. (**h**) Evaluation of the recombinant Siglec-10 antibody (clone 68A11A1) and anti-CD24 antibody in enhancing macrophage-mediated phagocytosis of multiple PDAC cell lines (AsPC-1, MIA PaCa-2, and PANC-1). Phagocytosis was normalized to the isotype control for each cell line and conducted using macrophages derived from monocytes of 5–8 healthy donors. Each symbol represents an individual donor; statistical significance was assessed using ratio paired t-tests compared to isotype control. Means with SEM are shown. (**i**) Triple co-culture assay involving cancer-associated fibroblasts (CAFs), PANC-1 PDAC cells, and monocyte-derived macrophages, showing phagocytosis kinetics, AUC quantification, and representative images. Statistical significance assessed using paired t-tests. (**j-k**) Flow cytometry analysis of CellTrace Violet (CTV)-labeled human CD8^+^ T cells co-cultured with human monocytes ± anti-Siglec-10 antibody in the presence of anti-CD3/CD28 beads for 5 days. (**j**) T cell proliferation; (**k**) granzyme B expression. ANOVA with post hoc comparisons.

To generate mAbs targeting Siglec-10, mice were immunized with recombinant human Siglec-10 protein, and serum antibody titers were measured using indirect ELISA. High reactivity to Siglec-10 was observed in several mice, which were subsequently selected for hybridoma generation. Over 200 hybridoma clones were initially screened for antibody production using binding specificities by ELISA against Siglec-10 and Siglec-5 (a control). Sixteen clones (sequences in **supplementary Table 1**) were selected for subcloning. Antibodies from these clones were further characterized by ELISA and flow cytometry-based immune assays. Several antibodies exhibited high affinity for Siglec-10 in binding ELISA (**Fig. 6b**). To confirm the binding specificity of these antibodies to Siglec-10, CHO-K1 cells expressing Siglec-10 or Siglec-5 (as a control) were incubated with hybridoma supernatants and stained with an Alexa Fluor 488-conjugated secondary antibody. The antibodies showed significant binding to Siglec-10-expressing cells, but not to Siglec-5 controls (**Fig. 6c**).

The clones were then evaluated for their ability to enhance macrophage-mediated phagocytosis of PDAC cells using the same assay described in Fig. 5, compared against an isotype control, an anti-CD24 antibody, and a commercially available anti-Siglec-10 antibody (**Fig. 6d**). Several clones significantly improved the capacity of macrophages to phagocytose PDAC cells (**Fig. 6d**). The top five clones were further tested using macrophages derived from monocytes of multiple donors to assess reproducibility. Among these, one clone, 68A11A1, demonstrated superior performance and was selected for recombinant expression (**Fig. 6e–f**). The recombinant 68A11A1 antibody (IgG1 isotype) showed high specificity for Siglec-10 without cross-reactivity to Siglec-5 or other irrelevant His-tagged proteins (**Fig. 6g**). Notably, this recombinant antibody significantly enhanced macrophage-mediated phagocytosis across multiple human PDAC cell lines compared to the anti-CD24 antibody, using macrophages derived from several donors (**Fig. 6h**). It also enhanced phagocytosis by monocyte-derived macrophages against PDAC PANC-1 cells even in the presence of cancer-associated fibroblasts (CAFs) in tri-culture assays with monocyte-derived macrophages, PDAC cells, and CAFs (**Fig. 6i**), and did so without altering the levels of ITGA3, ITGB1, or Siglec-10 ligands on PDAC cells (**Supplementary Fig. 6**).

In addition, we assessed the ability of this antibody to overcome myeloid cell-mediated suppression of T cell responses. In a monocyte–T cell suppression assay, Siglec-10 blockade increased CD8+ T cell proliferation (**Fig. 6j**) and granzyme B expression (**Fig. 6k**) compared to the isotype control. Together, these results demonstrate the successful generation and characterization of monoclonal antibodies with high affinity and specificity for Siglec-10. These antibodies not only enhanced macrophage-mediated phagocytosis of PDAC cells but also prevented myeloid cell-mediated inhibition of T cell proliferation and activation *in vitro*, underscoring the critical role of Siglec-10 in PDAC immune evasion and its potential as a therapeutic target.

### Disrupting Siglec-10 interactions enhances macrophage-mediated phagocytosis of PDAC cells and induces human macrophage activation in vivo

Since mice lack Siglec-10 and instead express its homolog Siglec-G, testing a Siglec-10 antibody in wild-type mice is not feasible. To overcome this limitation, we employed two complementary animal models. First, we used a xenograft model: immunodeficient NSG mice engrafted with AsPC-1 human PDAC cells and human monocyte-derived macrophages. Mice were treated with either an isotype control antibody or the Siglec-10-specific antibody clone 68A11A1 (**Fig. 7a**). Targeting Siglec-10 significantly reduced PDAC tumor growth *in vivo* (**Fig. 7b**).

**Figure 7.**
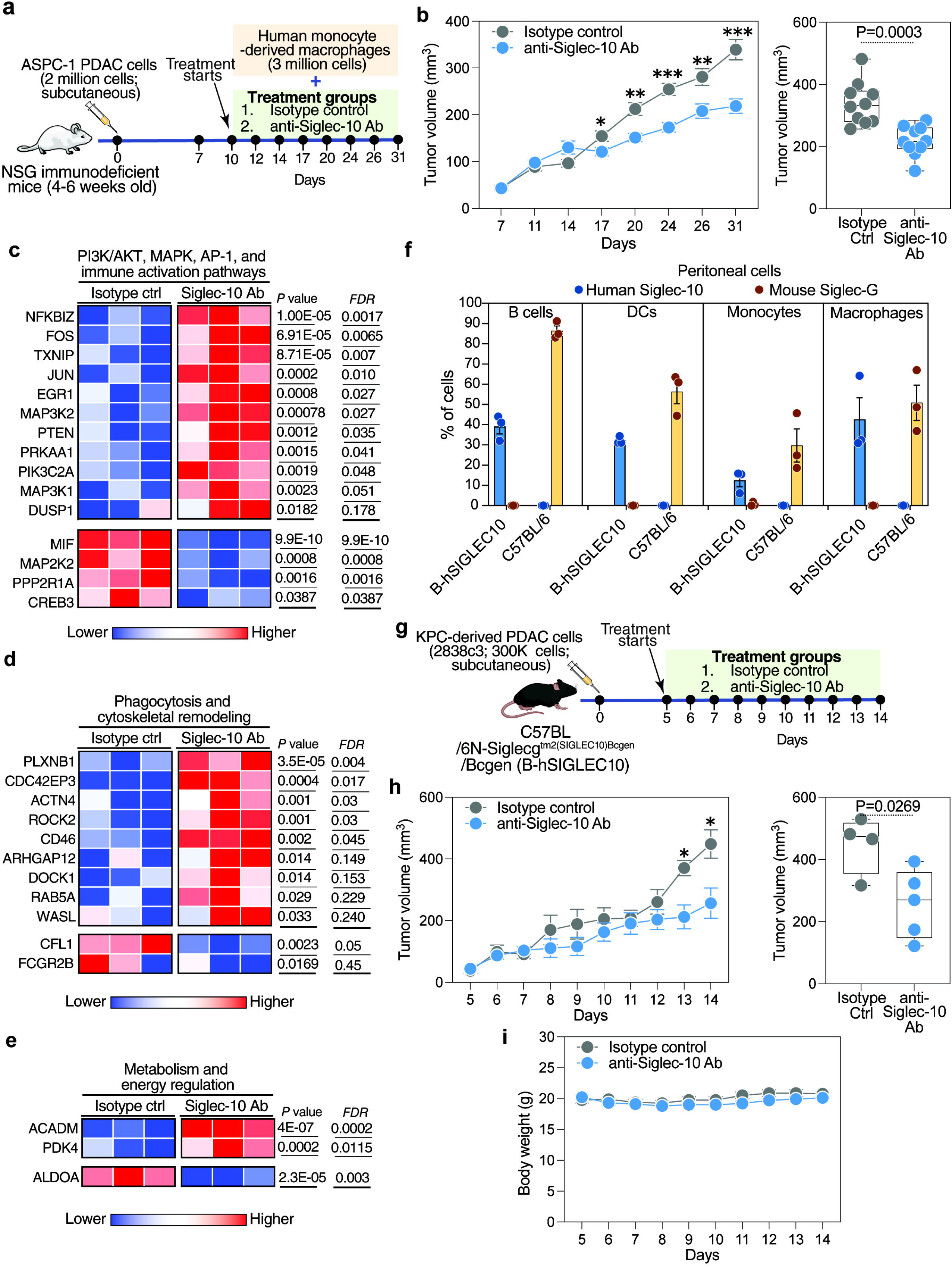
Blocking Siglec-10 interactions enhances macrophage-mediated phagocytosis and reduces PDAC tumor growth *in vivo*. (**a**) Experimental design for subcutaneous tumor development and Siglec-10 antibody treatment. AsPC-1 PDAC cells were subcutaneously injected into immunodeficient NSG mice. After 7 days, when tumors reached ∼40 mm³, mice were randomized into treatment groups. On the indicated days, ∼3 × 10 monocyte-derived macrophages were intravenously injected along with 200 µg of either isotype control or anti-Siglec-10 antibody. Tumor size was measured on specified days using a vernier caliper. (b) Tumor volume over time (left) and at the last time point (right) in mice treated with isotype control or anti-Siglec-10 antibody. n = 10 mice per group. Means with SEM are shown for the longitudinal analysis. For the last time point analysis, box and whiskers represent the minimum to maximum range, with all individual data points shown. Mann–Whitney U test. (**c–e**) Heatmaps showing differential gene expression in FACS-sorted human macrophages from tumors of NSG mice treated with anti-Siglec-10 antibody, highlighting: (c) immune activation pathways, (d) phagocytosis-related genes, and (e) metabolism-related genes. Red indicates higher expression; blue indicates lower expression. (**f**) Flow cytometry analysis of peritoneal cells from wild-type B6 mice and transgenic B-hSIGLEC10 mice. B6 mice express Siglec-G but not Siglec-10, whereas B-hSIGLEC10 mice express Siglec-10 but not Siglec-G on myeloid cells. Means with SEM are shown. (**g**) Experimental design for tumor challenge in B-hSIGLEC10 transgenic mice. KPC-derived murine PDAC cells (2838c3) were subcutaneously injected. After 5 days, mice were randomized and treated intravenously with either isotype control or anti– Siglec-10 antibody. (**h**) Tumor volume in B-hSIGLEC10 mice treated with isotype control or anti–Siglec-10 antibody. Tumor volume over time (left) and at the last time point (right) in mice treated with isotype control or anti-Siglec-10 antibody. Means with SEM are shown for the longitudinal analysis. N=4-5 per group. For the last time point analysis, box and whiskers represent the minimum to maximum range, with all individual data points shown. Unpaired t test. (**i**) Mouse body weight during treatment.

At the end of the experiment, tumors were harvested, and human macrophages were isolated by FACS (**Supplementary Fig. 7**). Transcriptomic analysis of these isolated macrophages by RNA sequencing revealed significant gene expression changes consistent with enhanced activation and phagocytic capacity upon Siglec-10 blockade (**Fig. 7c–e**). Specifically, macrophages from Siglec-10 antibody–treated mice exhibited enrichment of PI3K/AKT, MAPK, AP-1, and NF-κB signaling pathways, consistent with known Siglec-mediated signaling mechanisms described in previous studies (20,43–46). Upregulated genes included PIK3C2A, PTEN, and PRKAA1, important for PI3K/AKT signaling and macrophage metabolic reprogramming. Genes such as MAP3K1, MAP3K2, JUN, and FOS supported activation of MAPK and AP-1 pathways, promoting a pro-inflammatory and phagocytic phenotype. Additionally, upregulation of genes related to cytoskeletal remodeling (e.g., RAB5A, DOCK1, WASL) and downregulation of inhibitory genes (e.g., FCGR2B, CFL1) further underscored enhanced phagocytic function. Furthermore, macrophages displayed evidence of metabolic reprogramming, with upregulation of ACADM and PDK4 (supporting fatty acid oxidation) and downregulation of ALDOA (indicating reduced glycolysis), consistent with a profile favoring sustained immune activation. Together, these data show that Siglec-10 blockade *in vivo* reprograms macrophages toward an activated, phagocytic, and metabolically adaptable state.

To complement these findings, we next employed a genetically modified human Siglec-10 transgenic immunocompetent mouse model. Human Siglec-10 knock-in (B-hSIGLEC10) C57BL/6 mice are engineered to knock out Siglec-G and express human Siglec-10 instead (**Fig. 7f, Supplementary Fig. 8**). We subcutaneously engrafted these mice with 2838c3 cells, a highly aggressive murine PDAC cell line derived from the well-established KPC (LSL-Kras^G12D/+^; LSL-Trp53^R172H/+^; Pdx1-Cre) model. The KPC model is a spontaneous, autochthonous mouse model that faithfully recapitulates the genetic landscape, desmoplastic stroma, immune evasion, and rapid progression characteristic of human PDAC. We then treated these mice with either the Siglec-10 antibody or a control (**Fig. 7g**). Siglec-10 targeting significantly reduced tumor volume in this model without signs of toxicity (**Fig. 7h–i**). Overall, these *in vivo* results validate that blocking Siglec-10 reprograms macrophages by activating well-established immune signaling pathways (PI3K/AKT, MAPK, AP-1, NF-κB), consistent with mechanisms previously defined in the literature (20,43–46). Together, these findings support Siglec-10 blockade as a promising immunotherapeutic approach for PDAC.

## DISCUSSION

In this study, we identified a novel immunological interaction between α3β1 integrin on PDAC cell surfaces and Siglec-10, a glyco-immune checkpoint receptor on macrophages, which facilitates immune evasion by PDAC cells. This previously unrecognized interaction can suppress macrophage-mediated immune surveillance, enabling tumor cells to escape immune clearance. Notably, we show that disrupting Siglec-10 interactions with monoclonal antibodies restores macrophage phagocytic capacity against PDAC cells. By tapping into glycan-based interventions, a largely underexplored avenue in cancer immunotherapy, our findings lay the foundation for innovative strategies that could enhance therapeutic outcomes for PDAC patients. With PDAC projected to account for approximately 67,440 new cases and 51,980 deaths in the United States in 2025 alone (47), this work addresses an urgent and critical clinical need.

In 2019, a pivotal study identified interactions between Siglec-10 and CD24 as a key “don’t eat me” signal in breast and ovarian cancers, mediating immune evasion through macrophage suppression (20). Our findings uncover a distinct mechanism in PDAC, where α3β1 integrin, rather than CD24, emerges as one of the primary ligands for Siglec-10. Indeed, our comparative analyses of CD24 expression across PDAC, breast, and ovarian cancer cells showed that PDAC cells exhibit low CD24 expression, and many PDAC cells that bind Siglec-10 do not express CD24. This supports the presence of alternative Siglec-10 ligands in PDAC and underscores the importance of identifying context-dependent ligands mediating Siglec-10–driven immune evasion.

Integrin α3β1, composed of ITGA3 and ITGB1 subunits, is a well-characterized cell surface receptor known for roles in adhesion, migration, and metastasis (48–53). In PDAC, ITGA3 and ITGB1 are often overexpressed, contributing to epithelial-mesenchymal transition, invasion, and poor prognosis (54–60). Our study, for the first time, suggests a previously unrecognized immune regulatory function for α3β1 integrin, showing that its sialylated form acts as a critical ligand for Siglec-10, likely transmitting immunosuppressive signals that facilitate immune evasion. Our CRISPR-Cas9 knockout experiments of ITGA3 and depletion experiments of both ITGA3 and ITGB1 confirmed that loss of these integrins significantly increases susceptibility to macrophage-mediated phagocytosis. However, additional studies are needed to understand the exact glycomic modulation of these integrin interactions, as they are heavily glycosylated with multiple *N*-linked glycans and, in some contexts, may also carry *O*-linked glycans (61–66). This could be addressed through comprehensive glycoproteomic studies and genetic modulation of specific sialyltransferases and other glycosyltransferases that control the addition of Siglec-10 ligands to glycoproteins. These future investigations can broaden our understanding of α3β1 integrin, expanding its role to include immune modulatory properties within the tumor microenvironment.

Current immunotherapy efforts focusing on targeting proteins and nucleic acids have revolutionized cancer treatment. However, these approaches remain insufficient to fully realize the potential of immunotherapy for many patients, including those with PDAC (67,68). Targeting glycomic interactions between glycans and glyco-immune checkpoints such as Siglec-10 represents a novel and complementary approach to enhance the efficacy of cancer immunotherapy. Our findings underscore the therapeutic promise of disrupting Siglec-10 interactions as a strategy to restore macrophage function and enhance anti-PDAC immunity. Unlike traditional immune checkpoint inhibitors that primarily act on T cells (e.g., PD-1/PD-L1 blockers), Siglec-10 targeting directly reactivates myeloid cells like macrophages, which play a pivotal role within the tumor microenvironment. This creates opportunities for combining Siglec-10 targeting with other therapies. For example, combining it with therapies that target T cell checkpoints could further amplify anti-tumor immunity. Additionally, Siglec-10 targeting can be integrated with therapies designed to enhance macrophage-mediated phagocytosis, such as CD47-blocking antibodies, which are already in clinical development. CD47 inhibits phagocytosis by interacting with macrophage SIRPα (69–71). Since Siglec-10 and CD47 regulate phagocytosis through distinct pathways, targeting both simultaneously could produce additive or synergistic effects. Together, Siglec-targeting therapies, when combined with immune checkpoint inhibitors acting on T cells or macrophages, have the potential to significantly expand the arsenal of immunotherapeutic tools available for cancer treatment.

In our study, monoclonal antibodies against Siglec-10 restored macrophage phagocytic capacity *in vitro* and reduced tumor growth *in vivo*, providing proof of concept for targeting this glyco-immune checkpoint. However, targeting Siglecs on myeloid cells could have broader implications for immunity by activating antigen-presenting cells and potentially enhancing downstream T cell responses indirectly (72–74). While we focused on the role of Siglec-10 interactions in modulating phagocytosis, the broader impact of disrupting these interactions on overall anti-tumor immunity, including effects on T cells via modulation of MDSCs and DCs, warrants further investigation.

While our study provides significant insights into the role of Siglec-10 interactions in PDAC immune evasion, it is not without limitations. First, a deeper mechanistic investigation into the exact glycomic structures governing the binding between integrin α3β1 and Siglec-10 is needed, which could be accomplished by genetic manipulation of key sialyltransferases and other glycosyltransferases involved in adding Siglec-10 ligands to glycoproteins. Similarly, future studies could benefit from direct genetic manipulation of Siglec-10 in primary myeloid cells. Second, while we identified α3β1 integrin as a critical ligand for Siglec-10 in PDAC, it is likely that additional ligands exist in other cancer types, and these remain to be explored. Third, potential off-target effects and long-term consequences of Siglec-10 blockade, such as non-specific inflammation or autoimmunity, require further investigation in more complex and clinically relevant models. If toxicity is observed when blocking Siglecs using monospecific antibodies, alternative approaches may be required. Bispecific or trispecific antibodies targeting both Siglec-10 and its ligands, such as the α3β1 integrin identified in this study, could represent a promising strategy. Such approaches could enhance immune responses by disrupting Siglec-10-mediated immune suppression while minimizing the risk of non-specific inflammatory effects. Fourth, our study primarily focused on establishing the functional role of the Siglec-10 axis as an immune checkpoint that inhibits phagocytosis of PDAC cells using peripheral monocyte-derived macrophages to provide a controlled, reproducible system. However, the impact of macrophage polarization within the TME is highly relevant. Our single-cell analyses show that Siglec-10 is highly expressed on TAMs from human PDAC tumors and is associated with immunosuppressive transcriptional programs. This suggests that the Siglec-10 axis may be active even within immunosuppressive TAMs. Further studies directly examining reprogramming of TAMs isolated from PDAC patients would be highly valuable.

Despite these limitations, our findings represent a substantial advancement in understanding glyco-immune interactions in PDAC biology. By identifying α3β1 integrin as a novel ligand for Siglec-10 and showing the therapeutic potential of disrupting Siglec-10 interactions, we provide a new framework for targeting immune evasion in PDAC.

## SUPPLEMENTARY FIGURE LEGENDS

**Supplementary Figure 1. Pancreatic cancer is associated with high expression of sialyltransferases involved in producing Siglec-10 ligands. (a)** A schematic representation of the glycan ligands of Siglec-10 on *N*-linked and *O*-linked glycans, along with the enzymes involved in their formation. **(b–c)** Expression levels of sialyltransferase enzymes responsible for Siglec-10 ligand formation compared between normal tissues and PAAD tissues in the TCGA dataset. Unpaired t tests. *P < 0.05.

**Supplementary Figure 2. Monocyte-derived macrophages express Siglec-10.** A flow cytometry-based analysis of monocytes and macrophages derived from monocytes for Siglec-10 expression.

**Supplementary Figure 3. ITGA3 and ITGAB1 highly expresses in PDAC cells.** Several PDAC cells were examined for their expression of (a) ITGA3 and (b) ITGB1 (x-axis) by flow cytometry.

**Supplementary Figure 4. The expression of Siglec-10 and Siglec-10 ligand forming enzymes associated with the expression of ITGA3 and ITGB1.** Spearman r correlation analysis in TCGA of (a) Siglec-10 with ITGA3, (b) ITGA3 with Siglec-10 ligand forming enzymes (ST3GAL1 and B3GNT3), (c) Siglec-10 with ITGB1, and (d) ITGB1 with Siglec-10 ligand forming enzymes (ST3GAL1 and B3GNT3).

**Supplementary Figure 5. ITGB1-expressing PDAC cells evade macrophage-mediated phagocytosis.** Columns coated with anti-ITGB1 antibodies were used to isolate ITGB1^low^ cells (cells passing through the columns without binding). Cells passing through uncoated columns (expressing high levels of ITGB1) were used as controls. **(a)** Schematic representation of the experimental setup to test the phagocytic capacity of monocyte-derived macrophages targeting control cells (enriched with ITGB1^hi^ cells) or ITGB1^low^ cells (n = 5–6). **(b-c)** Phagocytosis of control and ITGB1^low^ MIA PaCa-2 (f) or PANC1 (g) PDAC cells by macrophages from different donors. Top: Representative images showing phagocytosis (increased red indicates higher phagocytosis). Bottom left: Live imaging results (each symbol represents data from one donor). Bottom right: AUC data compiled from multiple donors. Statistical significance was assessed using ratio paired t-tests.

**Supplementary Figure 6. Targeting Siglec-10 using anti–Siglec-10 antibodies does not impact the levels of ITGA3, ITGB1, or Siglec-10 ligands on PDAC cells.** Percentage and MFI of (**a–b**) ITGA3, (**c–d**) ITGB1, and (**e–f**) Siglec-10 ligands measured by flow cytometry. ANOVA tests corrected with Benjamini, Krieger, and Yekutieli method. Means and SEM are shown.

**Supplementary Figure 7. Gating strategy for sorting the human tumor associated macrophages from mouse subcutaneous tumor.** Single cell suspension from mouse subcutaneous tumor was used to obtain human monocytes derived macrophages injected 12hr intravenously for RNA seq analysis. Mouse BV711-CD45^-^ and Human APC-CD45^+^ cells were sorted for RNA seq analysis.

**Supplementary Figure 8. B-hSIGLEC10 mice express human Siglec-10 on their myeloid cells. (a–b)** Gating strategy for cells from (**a**) B-hSIGLEC10 mice and (**b**) C57BL/6 mice. (**c**) Representative example of the expression of human Siglec-10 and mouse Siglec-G on B cells, monocytes, macrophages, and DCs from both mouse models.

## SUPPLEMENTARY TABLES

**Supplementary Table 1.** Sequences of the monoclonal antibodies against Siglec-10

## ACKNOWLEDGMENTS

M.A-M is supported by NIH grants (NIH R01AG092241, R01AI165079, R01AA029859, R01DK123733, and R01NS117458). M.A-M is also funded by the NIH-funded BEAT-HIV Martin Delaney Collaboratory to cure HIV-1 infection (1UM1Al126620). H.Y.T is supported by NCI R50CA221838, and the Wistar Shared Resources are supported by P30 CA010815.

## AUTHOR CONTRIBUTIONS

M.A.-M conceived and designed the study. P.S carried out the majority of the experiments. G.M, S.M.S.I, S.A.B, and A.P.B performed the animal experiments. R.S. supervised and designed the animal experiments. L.M.S. and J.F.H. supervised and helped perform the CRISPR-Cas9 experiments. J.C and H.-Y.T conducted the SPR and mass spectrometry analyses. K.M. consulted on the generation of the antibodies. H.T and R.Z interpreted the data. P.S and M.A.-M wrote the manuscript, and all authors edited it.

