## Supplementary Materials for "Targeting Siglec-10/α3β1 Integrin Interactions Enhances Macrophage-Mediated Phagocytosis of Pancreatic Cancer"

### Supplementary Figure 1

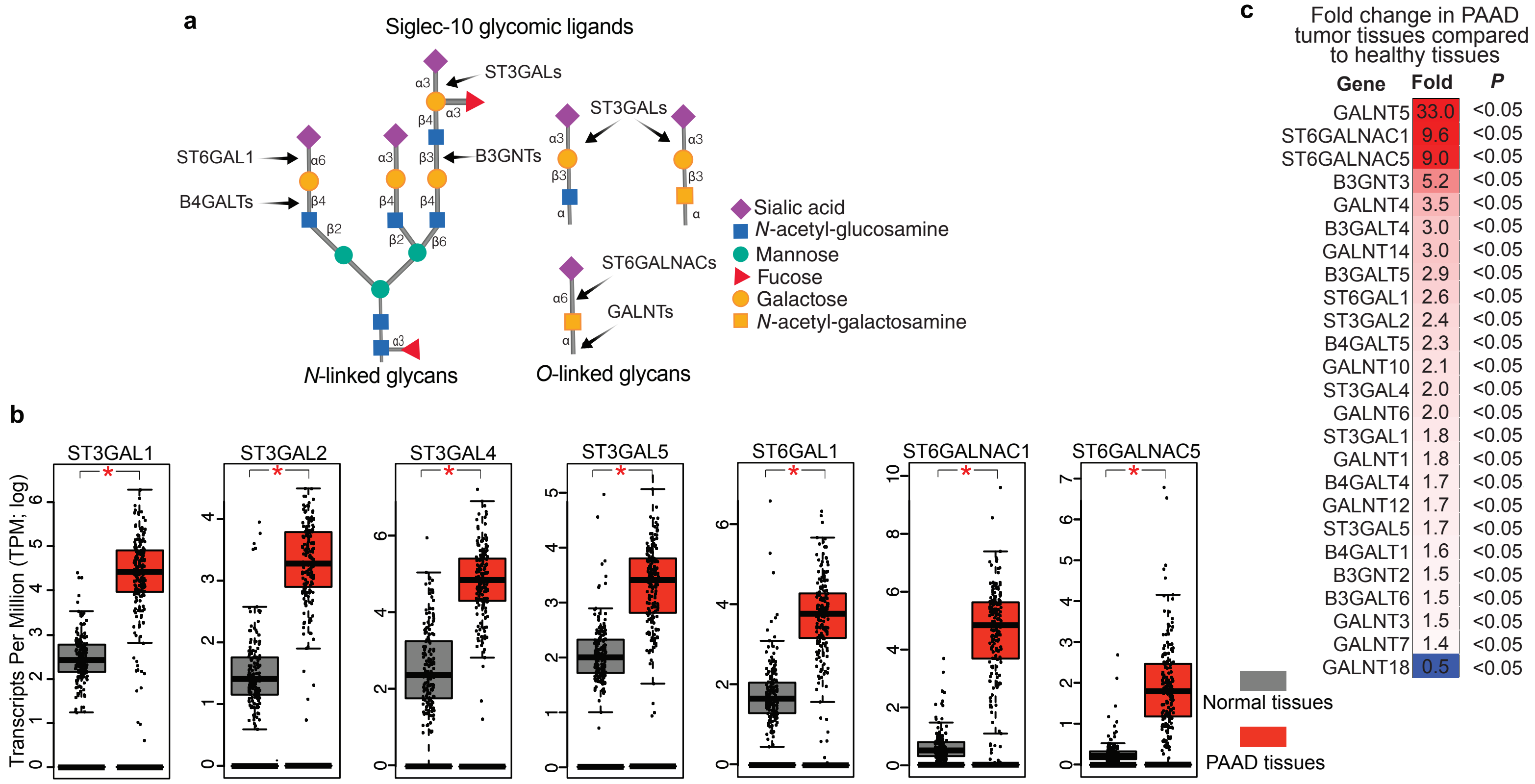

**Supplementary Figure 2**

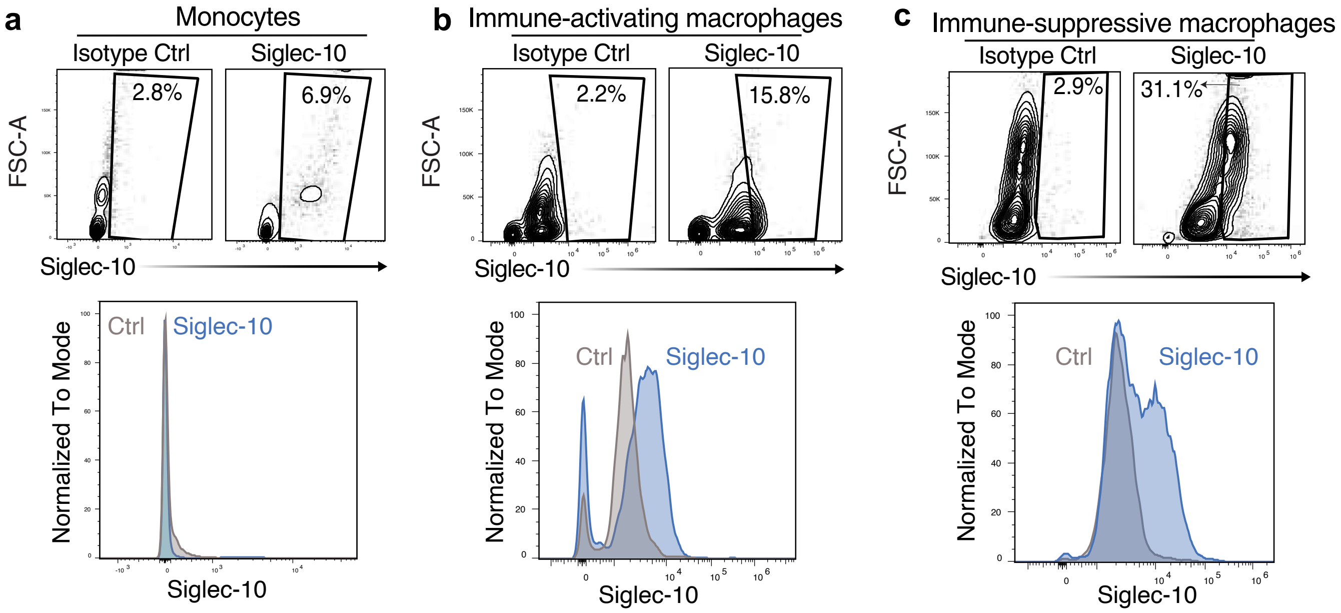

### Supplementary Figure 3

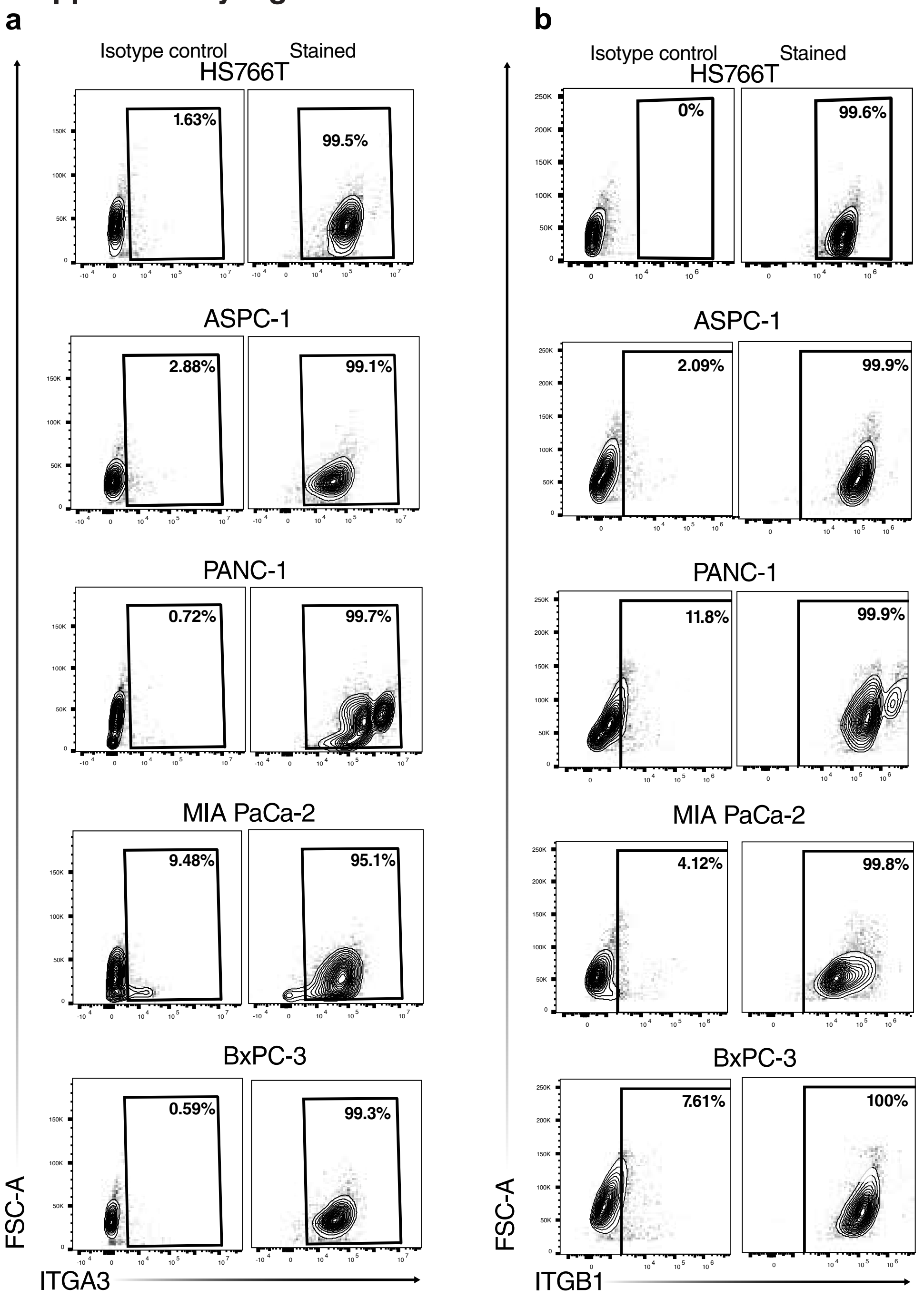

### Supplementary Figure 4

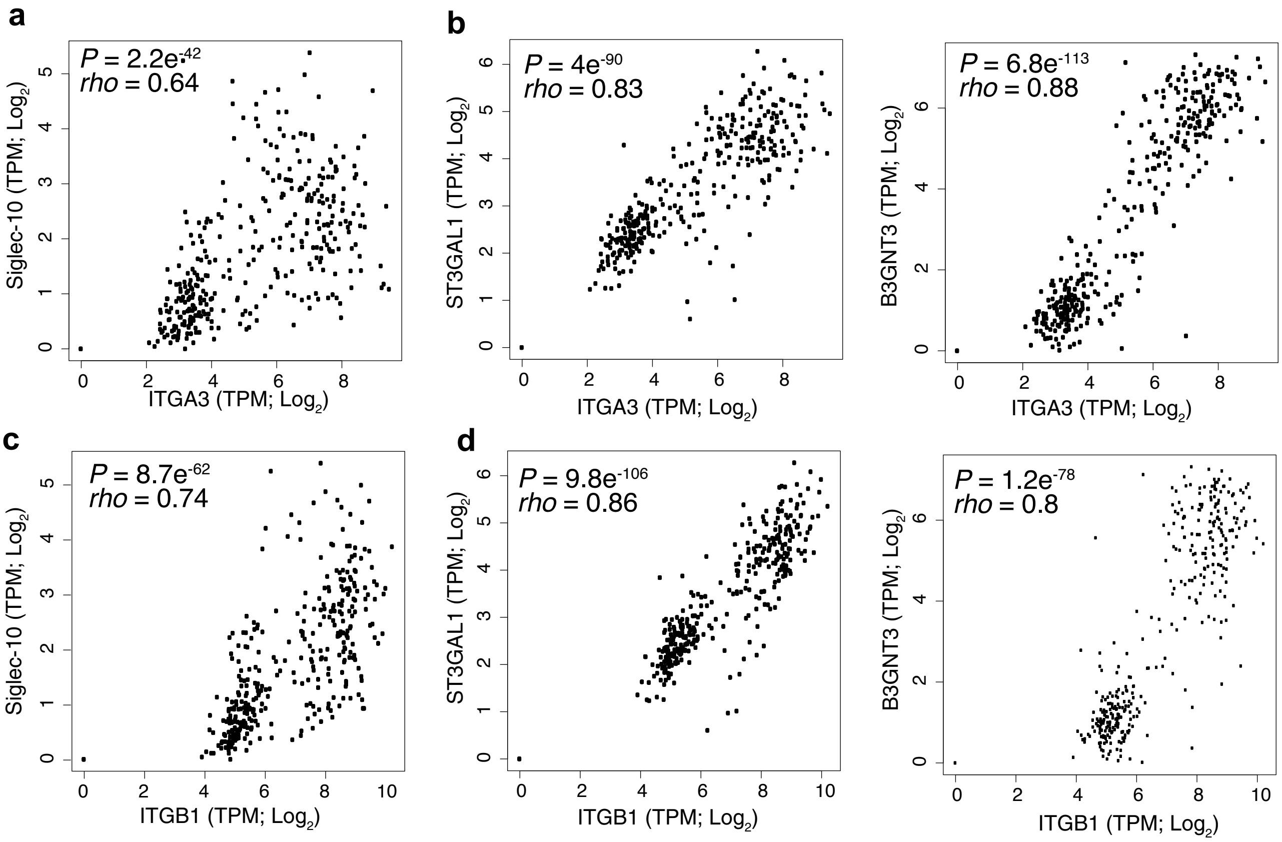

#### Supplementary Figure 5

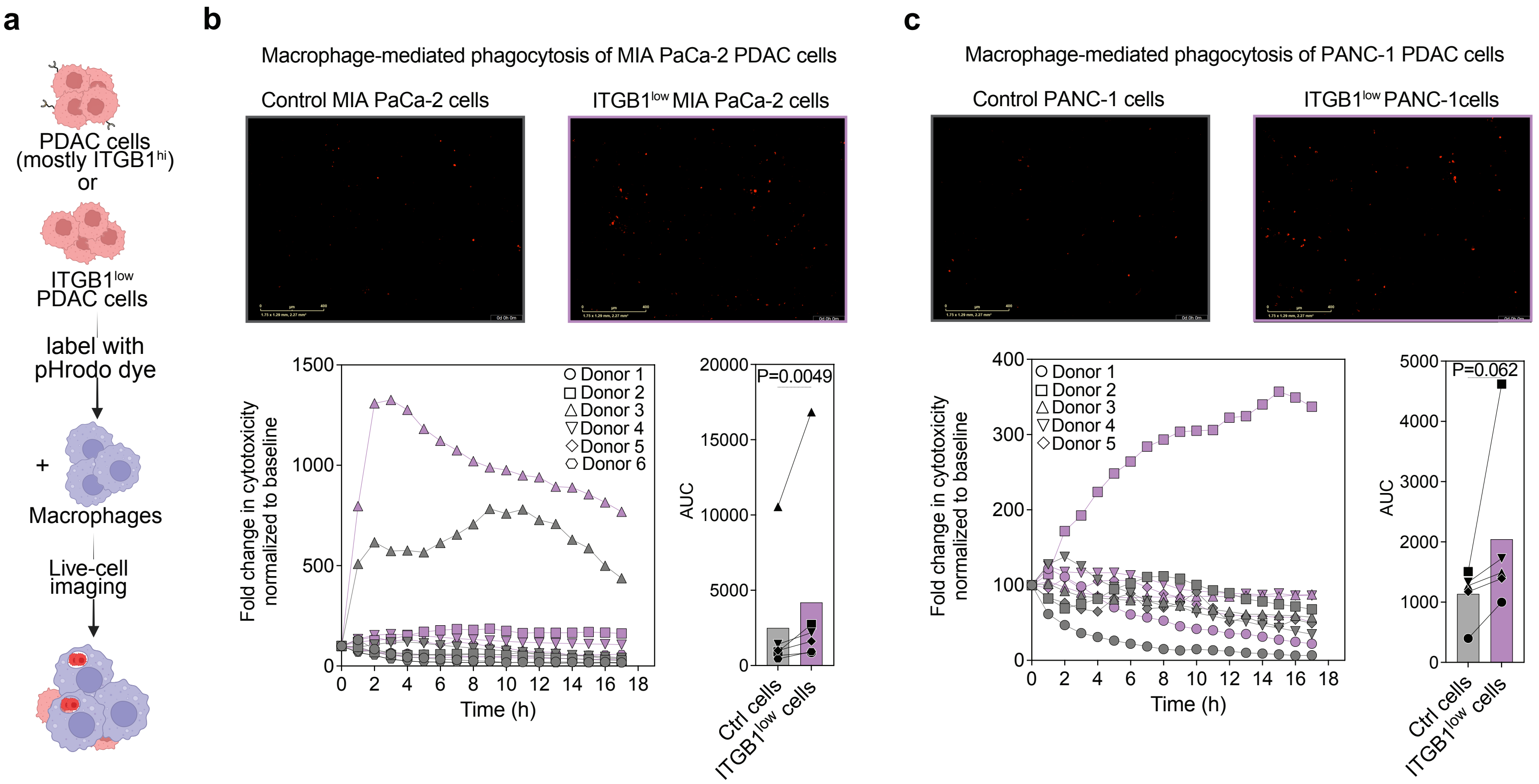

#### Supplementary Figure 6

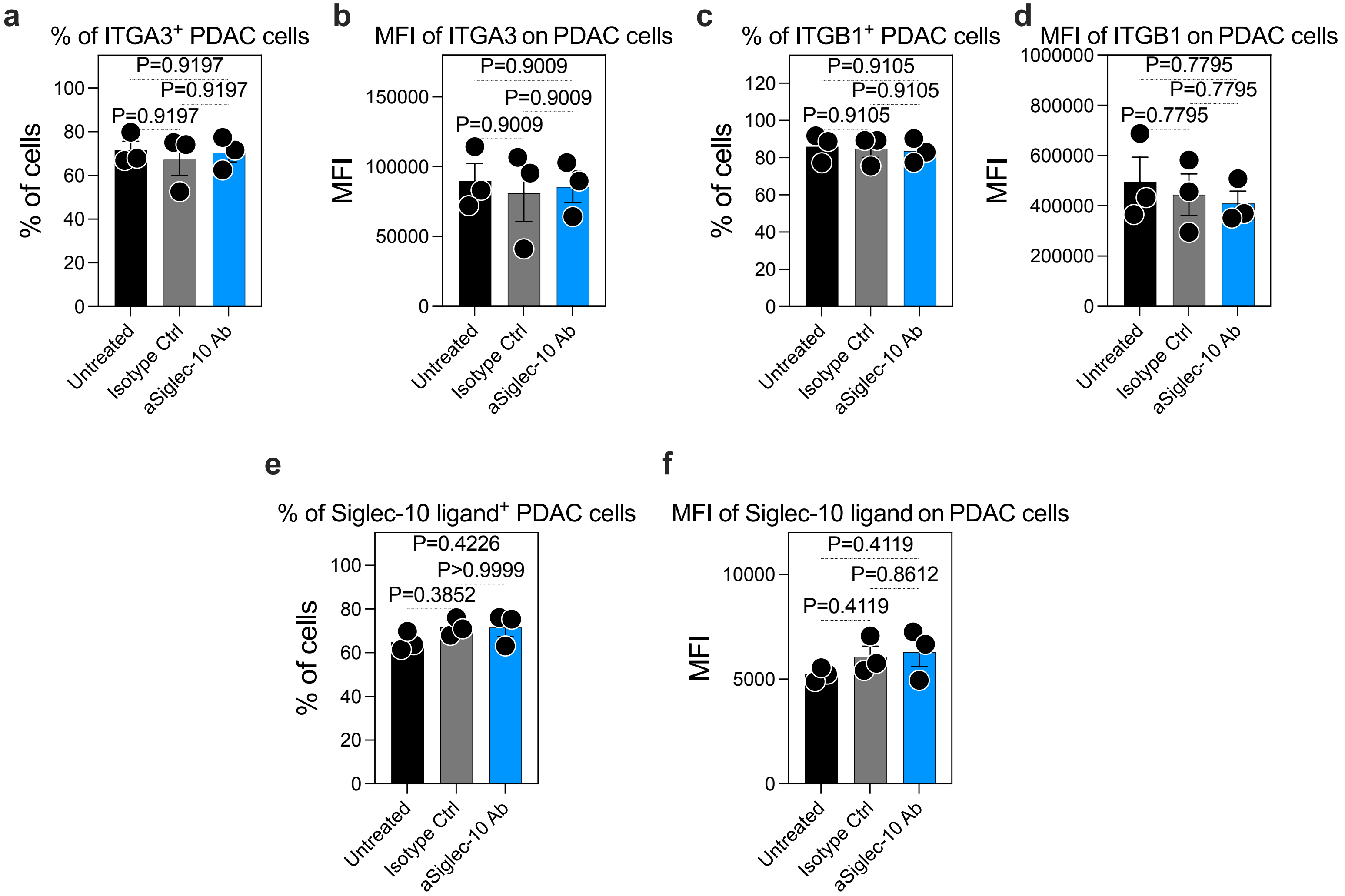

#### Supplementary Figure 7

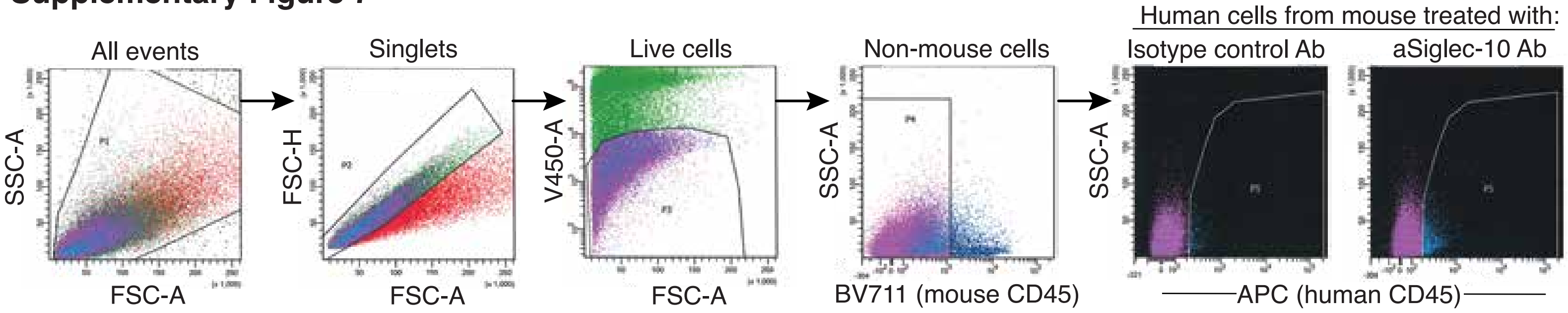

### Supplementary Figure 8

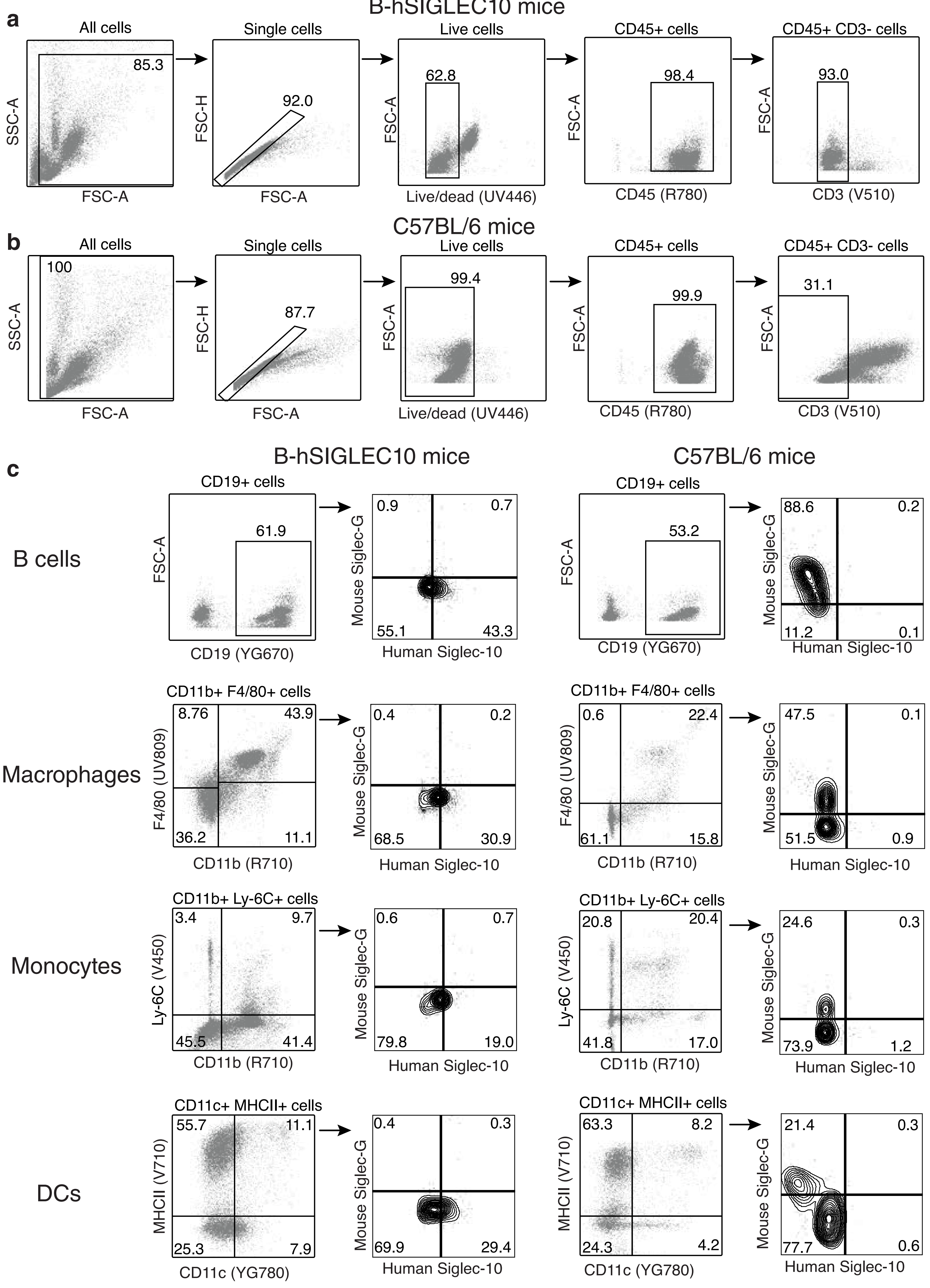

**Supplementary Table 1:** Sequences of Siglec-10 antibodies.

| Clone | Heavy chain: DNA sequence | Heavy chain: Amino acid sequence | Light chain: DNA sequence | Light chain: Amino acid sequence |
|---|---|---|---|---|
| **22F1A9** | ATGAACTTCGGGCTCAGCTTGATTTTCCTTGTCCTTGTTTTAAAAGGTGTCCAGTGTGAAGTGCAGCTGGTGGAGTCTGGGGGAGGCTTAGTGAAGCCTGGAGGGTCCCTGAAACTCTCCTGTGCAGCCTCTGGATTCACTTTCAGTAGCTATGCCATGTCTTGGGTTCGCCAGACTCCGGAAAAGAGGCTGGAGTGGGTCGCAACCATTAGTGATGGTGGTAGTTACACCTACTATCCAGACAATGTAAAGGGCCGATTCACCATCTCCAGAGACAATACCAAGAACAACCTGTACCTGCAAATGAGCCATCTGAAGTCTGAGGACACAGCCATGTTTTACTGTGCAAGAGATCTGTCCCCTCTCTACTATCGTGGGCCTATGGACTACTGGGGTCAAGGAACCTCAGTCACCGTCTCCTCA | MNFGLSLIFLVLVLKGVQCEVQLVESGGGLVKPGGSLKLSCAASGFTFSSYAMSWVRQTPEKRLEWVATISDGGSYTYYPDNVKGRFTISRDNTKNNLYLQMSHLKSEDTAMFYCARDLSPLYYRGPMDYWGQGTSVTVSS | ATGAGGGCTCCTGCACAGATTTTTGGCTTCTTGTTGCTCTTGTTTCCAGGTACCAGATGTGACATCCAGATGACCCAGTCTCCAGCTTCCTTATCTGCCTCTCTGGGAGAAAGAGTCAGTCTCACTTGTCGGGCAAGTCAGGACATTGGTAGTAGCTTAAACTGGCTTCAGCAGGAACCAGATGGAACTATTAAACGCCTGATCTACGCCACATCCAGTTTAGATTCTGGTGTCCCCAAAAGGTTCAGTGGCAGTAGGTCTGGGTCAGATTATTCTCTCACCATCAGCAGCCTTGAGTCTGAAGATTTTGTAGACTATTACTGTCTACAATATGCTAGTTCTCCATTCACGTTCGGCTCGGGGACAAAGTTGGAAATAAAA | MRAPAQIFGFLLLLFPGTRCDIQMTQSPASLSASLGERVSLTCRASQDIGSSLNWLQQEPDGTIKRLIYATSSLDSGVPKRFSGSRSGSDYSLTISSLESEDFVDYYCLQYASSPFTFGSGTKLEIK |
| **31E6H8** | ATGAAGTTGTGGCTGAACTGGATTTTCCTTGTAACACTTTTAAATGGTATCCAGTGTGAGGTGAAGCTGGTGGAGTCTGGAGGAGGCTTGGTACAGCCTGGGGGTTCTCTGAGTCTCTCCTGTGCAGCTTCTGGATTCACCTTCACTGATTACTACATGACCTGGGTCCGCCAGCCTCCAGGGAAGGCACTTGAGTGGTTGGGTTTTATTAGAAACAAAGCTAATGGTTACACAACAGAGTACAGTGCATCTGTGAAGGGTCGGTTCACCATCTCCAGAGATAATTCCCAAAGCATCCTCTCTCTTCAAATGAATGCCCTGAGCGCTGAGGACAGTGCCACTTATTACTGTGCAAGCCTTGTACTACGGGGGTTTGCTTACTGGGGCCAAGGGACTCTGGTCACTGTCTCTGCA | MKLWLNWIFLVTLLNGIQCEVKLVESGGGLVQPGGSLSLSCAASGFTFTDYYMTWVRQPPGKALEWLGFIRNKANGYTTEYSASVKGRFTISRDNSQSILSLQMNALSAEDSATYYCASLVLRGFAYWGQGTLVTVSA | ATGGAATCACAGACCCAGGTCCTCATGTTTCTTCTGCTCTGGGTATCTGGTGCCTGTGCAGACATTGTGATGACACAGTCTCCATCCTCCCTGGCTATGTCAGTAGGACAGAAGGTCACTATGAACTGCAAGTCCAGTCAGAGCCTTTTAAATAATAGCAATCAAAAGAACTATTTGGCCTGGTACCAGCAGAAACCAGGACAGTCTCCTAAACTTCTGGTATACTTTGCATCCACTAGGGAATCTGGGGTCCCTGATCGCTTCATAGGCAGTGGATCTGGGACAGATTTCACTCTTACCCTCAGCAGTGTGCAGGCTGAAGACCTGGCAGATTACTTCTGTCAACAACATTATATCACTCCGTACACGTTCGGAGGGGGGACCAAGCTGGAGATAAAA | MESQTQVLMFLLLWVSGACADIVMTQSPSSLAMSVGQKVTMNCKSSQSLLNNSNQKNYLAWYQQKPGQSPKLLVYFASTRESGVPDRFIGSGSGTDFTLTLSSVQAEDLADYFCQQHYITPYTFGGGTKLEIK |
| **35A3C10** | ATGAAGTTGTGGCTGAACTGGATTTTCCTTGTAACACTTTTAAATGGTCTCCAGTGTGAGGTGAAACTGGTGGAGTCTGGAGGAGGCTTGGTACAGCCTGGGGGTTCTCTGAGTCTCTCCTGTGCAGCTTCTGGATTCACCTTCACTGATTACTACATGAGTTGGGTCCGCCAGCCTCCAGGGAAGGCACTTGAGTGGTTGGGTTTTATTAGAAATAAAGTTAGTGGTTACACAACAGACTACAGTTCATCTGTGAAGGGTCGGTTCACCTTCTCCAGAGATAATTCCCAAAGCATCGTCTATCTTCATATGAATGCCCTGAGAGCTGAGGACAGTGCCACTTATTACTGTACAAGATATGTATCGCTATGGTTACTCGGGTACTTCGATGTCTGGGGCACAGGGACCACGGTCACCGTCTCCTCA | MKLWLNWIFLVTLLNGLQCEVKLVESGGGLVQPGGSLSLSCAASGFTFTDYYMSWVRQPPGKALEWLGFIRNKVSGYTTDYSSSVKGRFTFSRDNSQSIVYLHMNALRAEDSATYYCTRYVSLWLLGYFDVWGTGTTVTVSS | ATGAGGTTCCAGGTTCAGGTTCTGGGGCTCCTTCTGCTCTGGATATCAGTTCCAGGTGCCCAGTGTGATGTCCAGATAACCCAGTCTCCATCTTATCTTGCTGCATCTCCTGGAGAAACCATTACTATTAATTGCAGGGCAAGTAAGAGCATTAGCAAATATTTAGCCTGGTATCAAGAGAAACCTGGGAAAACTAATAAGCTTCTTATCTACTCTGGATCCACTTTGCAATCTGGAATTCCATCAAGGTTCAGTGGCAGTGGATCTGGTACAGATTTCACTCTCACCATCAGTAGCCTGGAGCCTGAAGATTTTGCAATGTATTACTGTCAACAGCATAATGAATACCCGCTCACATTCGGTGCTGGGACCAAGCTGGAGCTGAAA | MRFQVQVLGLLLLWISVPGAQCDVQITQSPSYLAASPGETITINCRASKSISKYLAWYQEKPGKTNKLLIYSGSTLQSGIPSRFSGSGSGTDFTLTISSLEPEDFAMYYCQQHNEYPLTFGAGTKLELK |
| **35B3F9** | ATGAAGTTGTGGCTGAACTGGATTTTCCTTGTAACACTTTTAAATGGTATCCAGTGTGAGGTGAAGCTGGTGGAGTCTGGAGGAGGCTTGGTACAGCCTGGGGGTTCTCTGAGGCTCTCCTGTGCAGCTTCTGGATTCACCTTCACTGATTACTACATGAGCTGGGTCCGCCAGCCCCCAGGGAAGGCACTTGAGTGGTTGGGTTTTATTAGAAATAAAGTTAGTGGTTACACAACAGACAACAGTGCATCTGTGAAGGGTCGGTTCACCATCTCCAGAGATGATTCCCAAAGCATCGTCTATCTTCAAATGAATGACCTGAGAGCTGAGGACAGTGCCACCTATTACTGTGCAAGATATGTTTCGCTTTGGTTACTCGGGTACTTCGATGTCTGGGGCACAGGGACCACGGTCACCGTCTCCTCA | MKLWLNWIFLVTLLNGIQCEVKLVESGGGLVQPGGSLRLSCAASGFTFTDYYMSWVRQPPGKALEWLGFIRNKVSGYTTDNSASVKGRFTISRDDSQSIVYLQMNDLRAEDSATYYCARYVSLWLLGYFDVWGTGTTVTVSS | ATGAGGTTCCAGGTTCAGGTTCTGGGGCTCCTTCTGCTCTGGATATCAGGTGCCCAGTGTGATGTCCAGATAACCCAGTCTCCATCTTATCTTGCTGCATCTCCTGGAGAAACCATTACTATTAATTGCAGGGCAAGTAAGAGCATTAGCAAATATTTAGCCTGGTATCAAGAGAAACCTGGGAAAACTAATAAGCTTCTTATCTACTCTGGATCCACTTTGCAATCTGGAATTCCATCAAGGTTCAGTGGCAGTGGATCTGGCACAGATTTCACTCTCACCATCAGTAGCCTGGAGCCTGAAGATTTTGCAATGTATTACTGTCAACAGCATAATGAATACCCACTCACATTCGGTACTGGGACCAAGCTGGAGCTGAAA | MRFQVQVLGLLLLWISGAQCDVQITQSPSYLAASPGETITINCRASKSISKYLAWYQEKPGKTNKLLIYSGSTLQSGIPSRFSGSGSGTDFTLTISSLEPEDFAMYYCQQHNEYPLTFGTGTKLELK |
| **35G6G1** | ATGAAGTTGTGGCTGAACTGGATTTTCCTTGTAACACTTTTAAATGGTATCCAGTGTGAGGTGAAGCTGGTGGAGTCTGGAGGAGGCTTGGTACAGCCTGGGGGTTCTCTGAGTCTCTCCTGTGCAGCTTCTGGATTCACCTTCGCTGATTACTACATGACCTGGGTCCGCCAGCCTCCAGGGAAGGCACTTGAGTGGTTGGGTTTTATTAGAAACAAAGCTAATGGTTTCACAACAGAGTACAGTGCATCTGTGAAGGGTCGGTTCACCATCTCCAGAGATAATTCCCAAAGCATCCTCTATCTTCAAATGAATGCCCTGAGAGCTGAGGACAGTGCCACTTATTACTGTGCAAGATATGGGTTACGGGGGTTTGCTTACTGGGGCCAAGGGACTCTGGTCACTGTCTCTGCA | MKLWLNWIFLVTLLNGIQCEVKLVESGGGLVQPGGSLSLSCAASGFTFADYYMTWVRQPPGKALEWLGFIRNKANGFTTEYSASVKGRFTISRDNSQSILYLQMNALRAEDSATYYCARYGLRGFAYWGQGTLVTVSA | ATGGAATCACAGACCCAGGTCCTCATGTTTCTTCTGCTCTGGGTATCTGGTGCCTGTGCAGACATTGTGATGACACAGTCTCCATCCTCCCTGGCTATGTCAGTAGGACAGAAGGTCACTTTGAACTGCAAGTCCAGTCAGAGCCTTTTAAATAGTAGCAATCAAAAGAACTATTTGGCCTGGTACCAGCAGAAACCAGGACAGTCTCCTAAACTTCTGGTATACTTTGCATCCACTAGGGAATCTGGGGTCCCTGATCGCTTCATAGGCAGTGGATCTGGGACAGATTTCACTCTTACCATCAGCAGTGTGCAGGCTGAAGACCTGGCAGATTACTTCTGTCACCAACATTATAGTACTCCATTCACGTTCGGCTCGGGGACAAAGTTGGAAATGAAA | MESQTQVLMFLLLWVSGACADIVMTQSPSSLAMSVGQKVTLNCKSSQSLLNSSNQKNYLAWYQQKPGQSPKLLVYFASTRESGVPDRFIGSGSGTDFTLTISSVQAEDLADYFCHQHYSTPFTFGSGTKLEMK |
| **36F7B3** | ATGAAGTTGTGGCTGAACTGGATTTTCCTTGTAACACTTTTAAATGGTATCCAGTGTGAGGTGAAGCTGGTGGAGTCTGGAGGAGGCTTGGTACAGCCTGGGGGTTCTCTGAGTCTCTCCTGTGCAGCTTCTGGATTCACCTTCACTGATTACTACATGACCTGGGTCCGCCAGCCTCCAGGGAAGGCACTTGAGTGGTTGTGTTTTATTAGAAACAAAGCTAATGGTTACACAACAGAGTACAGTGCATCTGTGAAGGGTCGGTTCACCATCTCCAGAGATAATTCTCAAAGCATCCTCTATCTTCAAGTGAATGCCCTGAGAGCTGAGGACAGTGCCACTTATTACTGTGCAACATATGTCCTACGTGGGCTTGCTTACTGGGGCCAAGGGACTCTGGTCACTGTCTCTGCA | MKLWLNWIFLVTLLNGIQCEVKLVESGGGLVQPGGSLSLSCAASGFTFTDYYMTWVRQPPGKALEWLCFIRNKANGYTTEYSASVKGRFTISRDNSQSILYLQVNALRAEDSATYYCATYVLRGLAYWGQGTLVTVSA | ATGGAATCACAGACCCAGGTCCTCATGTTTCTTCTGCTCTGGGTATCTGGTGCCTGTGCAGACATTGTGATGACACAGTCTCCATCCTCCCTGGCTATGTCAGTAGGACAGAAGGTCACTATGAGCTGCAAGTCCAGTCAGAGCCTTTTAAATAGTAGCAATCAAAAGAACTATTTGGCCTGGTACCAGCAGAAACCAGGACAGTCTCCTAAACTTCTGGTATACTTTGCATCCACTAGGGAATCTGGGGTCCCTGATCGCTTCATAGGCAGTGGATCTGGGACAAATTTCACTCTTACCATCAGCAGTGTGCAGGCTGAAGACCTGGCAGATTACTTCTGTCACCAACATTATAGCACTCCGCTCACGTTCGGTGCTGGGACCAAGCTGGAGCTGAAA | MESQTQVLMFLLLWVSGACADIVMTQSPSSLAMSVGQKVTMSCKSSQSLLNSSNQKNYLAWYQQKPGQSPKLLVYFASTRESGVPDRFIGSGSGTNFTLTISSVQAEDLADYFCHQHYSTPLTFGAGTKLELK |
| **38B2G5** | ATGGAAAGGCACTGGATCTTTCTCTCCCTGTTGTCAGTAATAGCAGGTGTCCACTCCCAGGTCCAGCTGCAGCAGTCTGGGGCTGAAGTGGCAAAACCTGGGGCCTCAGTGAAGCTGTCCTGCAAGGCTTCTGGCTACACCTTTACTAGCTACTGGATGCACTGGGTAAAACAGAGGCCTGGACAGGGTCTGGAATGGATTGGATACATTAATCCTAGCAATAATTATACTAAGTACAATCAGAAATTCAATGACAAGGCCACATTGACTGCAGACAAATCCTCCAGCACAGCCTACATGCAGCTGAGCAGCCTGACATATGAGGACTCTGCACTCTATTACTGTGCAAGAGTTTATTACTACGGTAATAGTCCTGCCTGGTTTGCTTACTGGGGCCAAGGGACTCTGGTCACTGTCTCTGCA | MERHWIFLSLLSVIAGVHSQVQLQQSGAEVAKPGASVKLSCKASGYTFTSYWMHWVKQRPGQGLEWIGYINPSNNYTKYNQKFNDKATLTADKSSSTAYMQLSSLTYEDSALYYCARVYYYGNSPAWFAYWGQGTLVTVSA | ATGGATTTTCAAGTGCAGATTTTCAGCTTCCTGCTAATCAGTGCTTCAGTCATAATGTCCAGAGGACAAATTGTTCTCTCCCAGTCTCCAGCAATCCTGTCTGCATCTCCAGGGGAGAAGGTCACAATGACTTGCAGGGCCAGCTCAAGTGTAAGTTACATGCACTGGTACCAGCAGAAGCCAGGATCCTCCCCCAAACCCTGGATTTATGCCACATCCAACCTGGCTTCTGGAGTCCCTGCTCGCTTCAGTGGCAGTGGGTCTGGGACCTCTTACTCTCTCACAATCAGCAGAGTGGAGGCTGAAGATGCTGCCACTTATTACTGCCAGCAGTGGACTAGTAACCCACCCACGTTCGGAGGGGGGACCAAGCTGGAAATAAAA | MDFQVQIFSFLLISASVIMSRGQIVLSQSPAILSASPGEKVTMTCRASSSVSYMHWYQQKPGSSPKPWIYATSNLASGVPARFSGSGSGTSYSLTISRVEAEDAATYYCQQWTSNPPTFGGGTKLEIK |
| **38D6A8** | ATGAAGTTGTGGCTGAACTGGATTTTCCTTGTAACACTTTTAAATGGTATCCAGTGTGAGGTGAAGCTGGTGGAGTCTGGAGGAGGCTTGGTACAGCCTGGGGGTTCTCTGAGTCTCTCCTGTGCAGCTTCTGGATTCACCTTCACTGATTACTACATGAGCTGGGTCCGCCAGCCTCCAGGGAAGGCACTTGAGTGGTTGGGTTTTATTAGAAACAAAGCTAGTGGTTACACAACAGAGTACAGTGCATCTGTGAAGGGTCGGTTCACCATCTCCAGAGATAATTCCCAAAACATCCTCTATCTTCAAATGAATGCCCTGAGAGCTGAGGACGGTGCCACTTATTACTGTGTAAGATATATATCGCTATGGTTACTCGGGTACTTCGATGTCTGGGGCACAGGGACCACGGTCACCGTCTCCTCA | MKLWLNWIFLVTLLNGIQCEVKLVESGGGLVQPGGSLSLSCAASGFTFTDYYMSWVRQPPGKALEWLGFIRNKASGYTTEYSASVKGRFTISRDNSQNILYLQMNALRAEDGATYYCVRYISLWLLGYFDVWGTGTTVTVSS | ATGAGGTTCCAGGTTCAGGTTCTGGGGCTCCTTCTGCTCTGGATATCAGGTGCCCAGTGTGATGTCCAGATAACCCAGTCTCCATCTTATCTTGCTGCATCTCCTGGAGAAACCATTACTATTAATTGCAGGGCAAGTAAGAGCATTAGCAAATATTTAGCCTGGTATCAAGAGAAACCTGGGAAAACTAATAAGCTTCTTATCTATTCTGGATCCACTTTGCAATCTGGAATTCCATCAAGGTTCAGTGGCAGTGGATCTGGTACAGATTTCACTCTCACCATCAGTAGCCTGGAGCCTGAAGATTTTGCAATGTATTACTGTCAACAGCATAATGATTATCCGCTCACATTCGGTGCTGGGACCAAGCTGGAGCTGAAA | MRFQVQVLGLLLLWISGAQCDVQITQSPSYLAASPGETITINCRASKSISKYLAWYQEKPGKTNKLLIYSGSTLQSGIPSRFSGSGSGTDFTLTISSLEPEDFAMYYCQQHNDYPLTFGAGTKLELK |
| **48D6F4** | ATGAAGTTGTGGCTGAACTGGATTTTCCTTGTAACACTTTTAAATGGTATCCAGTGTGAGGTGAGGCTGGTGGAGTCTGGAGGAGGCTTGGTACAGCCTGGGGGTTCTCTGCGTCTCTCCTGTGCAGCTTCTGGATTCACCTTCACTGATTACTACATGAGCTGGGTCCGCCAGCCTCCAGGGAAGGCACTTGAGTGGTTGGGTTTTATTAGAAACAAACCTAGTGGTTACACAACAGAATACAGTGCATCTGTGAAGGGTCGGTTCACCTTCTCCAGAGATAATTCCCAAAGCATCCTCTATCTTCAAATGAATGCCCTGAGAGCTGAGGACAGTGCCACTTACTACTGTGCAAGATATATATCGCTATGGTTACTCGGCTACCTCGATGTCTGGGGCACAGGGACCACGGTCACCGTCTCCTCA | MKLWLNWIFLVTLLNGIQCEVRLVESGGGLVQPGGSLRLSCAASGFTFTDYYMSWVRQPPGKALEWLGFIRNKPSGYTTEYSASVKGRFTFSRDNSQSILYLQMNALRAEDSATYYCARYISLWLLGYLDVWGTGTTVTVSS | ATGAGGTTCCAGGTTCAGGTTCTGGGGCTCCTTCTGCTCTGGATATCAGGTGCCCAGTGTGATGTCCAGATAACCCAGTCTCCATCTTATCTTGCTGCATCTCCTGGAGAAACCATTATTATTAATTGCAGGACAAGTAAGAGCATTAGCAAATATTTAGCCTGGTATCAAGAGAAACCTGGGAAAACTAATAACCTTCTTATCTACTCTGGATCCACTTTGCAATCTGGAATTCCATCAAGGTTCAGTGGCAGTGGATCTGGTACAGATTTCTCTCTCACCATCAGTAGCCTGGAGCCTGAAGATTTTGCAATCTATTACTGTCAACAACATAATGAATACCCGCTCACGTTCGGTGCTGGGACCAAGCTGGAGCTGAAT | MRFQVQVLGLLLLWISGAQCDVQITQSPSYLAASPGETIIINCRTSKSISKYLAWYQEKPGKTNNLLIYSGSTLQSGIPSRFSGSGSGTDFSLTISSLEPEDFAIYYCQQHNEYPLTFGAGTKLELN |
| **54C2F11** | ATGAAGTTGTGGCTGAACTGGATTTTCCTTGTAACACTTTTAAATGGTATCCAGTGTGAGGTGAGGCTGGTGGAGTCTGGAGGAGGCTTGGTACAGCCTGGGGGTTCTCTGCGTCTCTCCTGTGCAGCTTCTGGATTCACCTTCACTGATTACTACATGAGCTGGGTCCGCCAGCCTCCAGGGAAGGCACTTGAGTGGTTGGGTTTTATTAGAAACAAACCTAGTGGTTACACAACAGAGTACAGTGCATCTGTGAAGGGTCGGTTCACCATCTCCAGAGATAATTCCCAAAGCATCCTCTATCTTCAAATGAATGCCCTGAGAGCTGAGGACAGTGCCACTTACTACTGTGCAAGATATATATCGCTATGGTTACTCGGCTACCTCGATGTCTGGGGCACAGGGACCACGGTCACCGTCTCCTCA | MKLWLNWIFLVTLLNGIQCEVRLVESGGGLVQPGGSLRLSCAASGFTFTDYYMSWVRQPPGKALEWLGFIRNKPSGYTTEYSASVKGRFTISRDNSQSILYLQMNALRAEDSATYYCARYISLWLLGYLDVWGTGTTVTVSS | ATGAGGTTCCAGGTTCAGGTTCTGGGGCTCCTTCTGCTCTGGATATCAGGTGCCCAGTGTGATGTCCAGATAACCCAGTCTCCATCTTATCTTGCTGCATCTCCTGGAGAAACCATTATTATTAATTGCAGGACAAGTAAGAGCATTAGCAAATATTTAGCCTGGTATCAAGAGAAACCTGGGAAAACTAATAAGCTTCTTATCTACTCTGGATCCACTTTGCAATCTGGAATTCCATCAAGGTTCAGTGGCAGTGGATCTGGTACAGATTTCACTCTCACCATCAGTAGCCTGGAGCCTGAAGATTTTGCAATGTATTACTGTCAACAACATAATGAATACCCGCTCACGTTCGGTGCTGGGACCAAGCTGGAGCTGAAT | MRFQVQVLGLLLLWISGAQCDVQITQSPSYLAASPGETIIINCRTSKSISKYLAWYQEKPGKTNKLLIYSGSTLQSGIPSRFSGSGSGTDFTLTISSLEPEDFAMYYCQQHNEYPLTFGAGTKLELN |
| **54C3B11** | ATGAAGTTGTGGCTGAACTGGATTTTCCTTGTAACACTTTTAAATGGTATCCAGTGTGAGGTGAAGCTGGTGGAGTCTGGAGGAGGCTTGGTACAGCCTGGGGGTTCTCTGAGTCTCTCCTGTGCAGCTTCTGGATTCACCTTCACTGATTACTACATGACCTGGGTCCGCCAGCCTCCAGGGAAGGCACTTGAGTGGTTGTGTTTTATTAGAAACAAAGCTAATGGTTACACAACAGAGTACAGTGCATCTGTGAAGGGTCGGTTCACCATCTCCAGAGATAATTCTCAAAGCATCCTCTATCTTCAAGTGAATGCCCTGAGAGCTGAGGACAGTGCCACTTATTACTGTGCAACATATGTCCTACGTGGGCTTGCTTACTGGGGCCAAGGGACTCTGGTCACTGTCTCTGCA | MKLWLNWIFLVTLLNGIQCEVKLVESGGGLVQPGGSLSLSCAASGFTFTDYYMTWVRQPPGKALEWLCFIRNKANGYTTEYSASVKGRFTISRDNSQSILYLQVNALRAEDSATYYCATYVLRGLAYWGQGTLVTVSA | ATGGAATCACAGACCCAGGTCCTCATGTTTCTTCTGCTCTGGGTATCTGGTGCCTGTGCAGACATTGTGATGACACAGTCTCCATCCTCCCTGGCTATGTCAGTAGGACAGAAGGTCACTATGAGCTGCAAGTCCAGTCAGAGCCTTTTAAATAGTAGCAATCAAAAGAACTATTTGGCCTGGTACCAGCAGAAACCAGGACAGTCTCCTAAACTTCTGGTATACTTTGCATCCACTAGGGAATCTGGGGTCCCTGATCGCTTCATAGGCAGTGGATCTGGGACAAATTTCACTCTTACCATCAGCAGTGTGCAGGCTGAAGACCTGGCAGATTACTTCTGTCACCAACATTATAGCACTCCGCTCACGTTCGGTGCTGGGACCAAGCTGGAGCTGAAA | MESQTQVLMFLLLWVSGACADIVMTQSPSSLAMSVGQKVTMSCKSSQSLLNSSNQKNYLAWYQQKPGQSPKLLVYFASTRESGVPDRFIGSGSGTNFTLTISSVQAEDLADYFCHQHYSTPLTFGAGTKLELK |
| **56C3E6** | ATGAAGTTGTGGCTGAACTGGATTTTCCTTGTAACACTTTTAAATGGTATCCAGTGTGAGGTGAACCTGGTGGAGTCTGGAGGAGGCTTGGTACAGCCTGGGGATTCTCTGAGTCTCTCCTGTGCAACTTCTGGATTCACCTTCACTGATTACTACATGAGCTGGGTCCGCCAGCCTCCCGGGAAGGCACTTGAGTGGTTGGGTTTTATTAGAAACAAAGCTAATGGTTACACAACAGAGTACAGTGCATCTGTGAAGGGTCGGTTTACCATCTCCAGAGATAATTCCCAAAGCATCCTCTATCTTCAAATGAATGCCCTGAGACCTGAGGACAGTGCCACTTATTACTGTGCAAGCCCTGTACTACGGGGGTTTGCTTACTGGGGCCGAGGGACTCTGGTCACTGTCTCTGCA | MKLWLNWIFLVTLLNGIQCEVNLVESGGGLVQPGDSLSLSCATSGFTFTDYYMSWVRQPPGKALEWLGFIRNKANGYTTEYSASVKGRFTISRDNSQSILYLQMNALRPEDSATYYCASPVLRGFAYWGRGTLVTVSA | ATGAGTGTGCTCACTCAGGTCCTGGCGTTGCTGCTGCTGTGGCTTACAGGTGCCAGATGTGACATCCAGATGACTCAGTCTCCAGCCTCCCTATCTGCATCTGTGGGAGAAACTGTCACCATCACATGTCGAGCAAGTGGGAATATTCACAATTATTTAGCATGGTATCAGCAGAAACAGGGAAAATCTCCTCAGCTCCTGGTCTATAATGCAAAAACCTTAGCAGATGGTGTGCCATCAAGGTTCAGTGGCAGTGGATCAGGAACACAATATTCTCTCAAGATCAACAGCCTGCAGCCTGAAGATTTTGGGAGTTATTACTGTCAACATTTTTGGAGTACTCCATTCACGTTCGGCTCGGGGACAAAGTTGGAAATAAAA | MSVLTQVLALLLLWLTGARCDIQMTQSPASLSASVGETVTITCRASGNIHNYLAWYQQKQGKSPQLLVYNAKTLADGVPSRFSGSGSGTQYSLKINSLQPEDFGSYYCQHFWSTPFTFGSGTKLEIK |
| **66E8F8** | ATGAAGTTGTGGCTGAACTGGATTTTCCTTGTAACACTTTTAAATGGTATCCAGTGTGAGGTGAAGCTGGTGGAGTCTGGAGGAGGCTTGGTACAGCCTGGGGGTTCTCTGAGTCTCTCCTGTGCAGCTTCTGGATTCACCTTCACTGATTACTACATGAGCTGGGTCCGCCAGCCTCCAGGGAAGGCACTTGAGTGGTTGTGTTTTATTAGAAACAAAGCTAATGGTTACACAACAGAGTACAGTGCATCTGTGAAGGGTCGGTTCACCATCTCCAGAGATAACTCCCAAAGCATCCTCTATCTTCAAGTGAATGCCCTGAGAGCTGAGGACAGTGCCACTTATTACTGTGCAACATATGTCCTACGTGGGCTTGCTTACTGGGGCCAAGGGACTCTGGTCACTGTCTCTGCA | MKLWLNWIFLVTLLNGIQCEVKLVESGGGLVQPGGSLSLSCAASGFTFTDYYMSWVRQPPGKALEWLCFIRNKANGYTTEYSASVKGRFTISRDNSQSILYLQVNALRAEDSATYYCATYVLRGLAYWGQGTLVTVSA | ATGGAATCACAGACCCAGGTCCTCATGTTTCTTCTGCTCTGGGTATCTGGTGCCTGTGCAGACATTGTGATGACACAGTCTCCATCCTCCCTGGCTATGTCAGTAGGACAGAAGGTCACTATGAGCTGCAAATCCAGTCAGAGCCTTTTAAATAGTAGCAATCAAAAGAACTATTTGGCCTGGTACCAGCAGAAACCAGGACAGTCTCCTAAACTTCTGGTATACTTTGCATCCACTAGGGAATCTGGGGTCCCTGATCGCTTCATAGGCAGTGGATCTGGGACAGATTTCACTCTTACCATCAGCAGTGTGCAGGCTGAGGACCTGGCAGATTACTTCTGTCACCAACATTATAGCACTCCGCTCACGTTCGGTGCTGGGACCAAGCTGGAGCTGAAA | MESQTQVLMFLLLWVSGACADIVMTQSPSSLAMSVGQKVTMSCKSSQSLLNSSNQKNYLAWYQQKPGQSPKLLVYFASTRESGVPDRFIGSGSGTDFTLTISSVQAEDLADYFCHQHYSTPLTFGAGTKLELK |
| **68A11A1** | ATGAAGTTGTGGCTGAACTGGATTTTCCTTGTAACACTTTTAAATGGTTTCCAGTGTGAGGTGAAACTGGTGGAGTCTGGAGGAGGCTTGGTACAGCCTGGGGGTTCTCTGAGTCTCTCCTGTGCAGCTTCTGGATTCACCTTCACTGATTACTACATGAGTTGGGTCCGCCAGCCTCCAGGGAAGGCACTTGAGTGGTTGGGTTTTATTAGAAACAAAGTTAGTGGTTACACAACAGACTACAGTTCATCTGTGAAGGGTCGGTTCACCTTCTCCAGAGATAATTCCCAAAGCATCGTCTATCTTCATATGAATGCCCTGAGAGCTGAGGACAGTGCCACTTATTACTGTACAAGATATGTATCGCTTTGGTTACTCGGGTACTTCGATGTCTGGGGCACAGGGACCACGGTCACCGTCTCCTCA | MKLWLNWIFLVTLLNGFQCEVKLVESGGGLVQPGGSLSLSCAASGFTFTDYYMSWVRQPPGKALEWLGFIRNKVSGYTTDYSSSVKGRFTFSRDNSQSIVYLHMNALRAEDSATYYCTRYVSLWLLGYFDVWGTGTTVTVSS | ATGAGGTTCCAGGTTCAGGTTCTGGGGCTCCTTCTGCTCTGGATATCAGTTCCAGGTGCCCAGTGTGATGTCCAGCTAACCCAGTCTCCATCTTATCTTGCTGCATCTCCTGGAGAAACCATTACTATTAATTGCAGGACAAGTAAGAGCATTAGCAAATATTTAGCCTGGTATCAAGAGAAACCTGGGAAAACTAATAAGCTTCTTATCTACTCTGGATCCACTTTGCAATCTGGAATTCCATCAAGGTTCAGTGGCAGTGGATCTGGTACAGATTTCACTCTCACCATCAGTAGCCTGGAGCCTGAAGATTTTGCAATGTATTACTGTCAACAGCATAATGAATACCCGCTCACATTCGGTGCTGGGACCAAGCTGGAGCTGAAA | MRFQVQVLGLLLLWISVPGAQCDVQLTQSPSYLAASPGETITINCRTSKSISKYLAWYQEKPGKTNKLLIYSGSTLQSGIPSRFSGSGSGTDFTLTISSLEPEDFAMYYCQQHNEYPLTFGAGTKLELK |
| **68B1E2** | ATGAAGTTGTGGCTGAACTGGATTTTCCTTGTAACACTTTTAAATGGTATCCAGTGTGAGGTGAAGCTGGTGGAGTCTGGAGGAGGCTTGGTACAGCCTGGGGGTTCTCTGAGTCTCTCCTGCGCAGCTTCTGGATTCACCTTCACTGATTACTACATGAGCTGGGTCCGCCAGCCTCCAGGGAAGGCACTTGAGTGGTTGGGTTTTATTAGAAACAAAGCTAGTGGTTACACAACAGAGTACAATGCATCTGTGAAGGGTCGGTTCACCATCTCCAGAGATAATTCCCAAAACATCCTCTATCTTCAAATGAATGCCCTGAGAGCTGAGGACGGTGCCACTTATTACTGTGTAAGATATATATCGCTATGGTTACTCGGGTACTTCGATGTCTGGGGCACAGGGACCACGGTCACCGTCTCCTCA | MKLWLNWIFLVTLLNGIQCEVKLVESGGGLVQPGGSLSLSCAASGFTFTDYYMSWVRQPPGKALEWLGFIRNKASGYTTEYNASVKGRFTISRDNSQNILYLQMNALRAEDGATYYCVRYISLWLLGYFDVWGTGTTVTVSS | ATGAGGTTCCAGGTTCAGGTTCTGGGGCTCCTTCTGCTCTGGATATCAGGTGCCCAGTGTGATGTCCAGATAACCCAGTCTCCATCTTATCTTGCTGCATCTCCTGGAGAAACCATTACTATTAATTGCAGGGCAAGTAAGAGCATTAGCAAATATTTAGCCTGGTATCAAGAGAAACCTGGGAAAACTAATAAGCTTCTTATCTATTCTGGATCCACTTTGCAATCTGGAATTCCATCAAGGTTCAGTGGCAGTGGATCTGGTACAGATTTCACTCTCACCATCAGTAGCCTGGAGCCTGAAGATTTTGCAATGTATTACTGTCAACAGCATAATGATTATCCACTCACATTCGGTGCTGGGACCAAGCTGGAGCTGAAA | MRFQVQVLGLLLLWISGAQCDVQITQSPSYLAASPGETITINCRASKSISKYLAWYQEKPGKTNKLLIYSGSTLQSGIPSRFSGSGSGTDFTLTISSLEPEDFAMYYCQQHNDYPLTFGAGTKLELK |
| **73D4G2** | ATGAAGTTGTGGCTGAACTGGATTTTCCTTGTAACACTTTTAAATGGTATCCAGTGTGAGGTGAGGCTGGTGGAGTCTGGAGGAGGCTTGGTACAGCCTGGGGGTTCTCTGCGTCTCTCCTGTGCAGCTTCTGGATTCACCTTCACTGATTACTACATGAGCTGGGTCCGCCAGCCTCCAGGGAAGGCACTTGAGTGGTTGGGTTTTATTAGAAACAAACCTAGTGGTTACACAACAGAGTACAGTGCATCTGTGAAGGGTCGGTTCACCATCTCCAGAGATAATTCCCAAAGCATCCTCTATCTTCAAATGAATGCCCTGAGAGCTGAGGACAGTGCCACTTACTACTGTGCAAGATATATATCGTTATGGTTACTCGGCTACCTCGATGTCTGGGGCACAGGGACCACGGTCACCGTCTCCTCA | MKLWLNWIFLVTLLNGIQCEVRLVESGGGLVQPGGSLRLSCAASGFTFTDYYMSWVRQPPGKALEWLGFIRNKPSGYTTEYSASVKGRFTISRDNSQSILYLQMNALRAEDSATYYCARYISLWLLGYLDVWGTGTTVTVSS | ATGAGGTTCCAGGTTCAGGTTCTGGGGCTCCTTCTGCTCTGGATATCAGGTGCCCAGTGTGATGTCCAGATAACCCAGTCTCCATCTTATCTTGCTGCATCTCCTGGAGAAACCATTATTATTAATTGCAGGACAAGTAAGAGCATTAGCAAATATTTAGCCTGGTATCAAGAGAAACCTGGGAAAACTAATAAGCTTCTTATCTACTCTGGATCCACTTTGCAATCTGGAATTCCATCAAGGTTCAGTGGCAGTGGATCTGGTACAGATTTCACTCTCACCATCAGTAGCCTGGAGCCTGAAGATTTTGCAATGTATTACTGTCAACAACATAATGAATACCCGCTCACGTTCGGTGCTGGGACCAAGCTGGAGCTGAAA | MRFQVQVLGLLLLWISGAQCDVQITQSPSYLAASPGETIIINCRTSKSISKYLAWYQEKPGKTNKLLIYSGSTLQSGIPSRFSGSGSGTDFTLTISSLEPEDFAMYYCQQHNEYPLTFGAGTKLELK |
